# Mast Cells in the Developing Brain Determine Adult Sexual Behavior

**DOI:** 10.1101/205559

**Authors:** Kathryn M. Lenz, Lindsay A. Pickett, Christopher L. Wright, Katherine T. Davis, Anabel Galan, Margaret M. McCarthy

**Author notes:** **Correspondence to:** Kathryn M. Lenz, 1835 Neil Ave, Columbus OH, 43210, USA.

## Abstract

Sex differences in brain and behavior are programmed during development by gonadal hormones. We found that the immune system-derived mast cell is a primary target for the masculinizing hormone, estradiol. Male rats had more mast cells in the preoptic area (POA), a brain region essential for male copulatory behavior, during the critical period for sexual differentiation. Activating mast cells in females masculinized POA neuronal and microglial morphology and adult sex behavior, and inhibiting mast cells in males blunted masculinization. Estradiol increased mast cell number and caused mast cells to release histamine, which stimulated microglia to release prostaglandins and thereby induced male-typical synaptic patterning. Inducing an allergic reaction in pregnant dams increased mast cell number in the brains of female fetuses and masculinized neuronal and microglia morphology and adult copulatory behavior. These findings identify a novel non-neuronal origin of brain sex differences and non-steroidal source of variability in brain feminization.

## Introduction

Elucidating the neural underpinnings that regulate complex social and motivated behaviors remains one of the unmet goals of neuroscience. Mating is a tightly orchestrated hormonally driven behavior that is expressed differently in males and females as a result of sexual differentiation of the brain early in development. The preoptic area (POA) is an essential brain region for expression of male copulatory behavior and embedded therein are multiple sex differences in neuron, astrocyte and microglial number and morphology (*1*). Establishing the cellular mechanisms by which sex differences are established provides unique insight into the neural control of behavior.

In perinatal male rodents, testicular androgens are converted to estrogens by neuronal aromatase and estrogens then induce masculinization of brain and behavior (*2*). Treatment of newborn female rodents with testosterone or its active metabolite, estradiol, induces masculinization via a complete phenotypic switching of neuroanatomical and behavioral endpoints (*3*). We previously determined that the inflammatory signaling molecule, prostaglandin E2 (PGE2), is induced by estradiol in the neonatal POA and is both necessary and sufficient for the masculinization process (*4*). In males, microglia exhibited a morphology consistent with a higher degree of activation and production of inflammatory signaling molecules, including PGE2. The presence of activated (ameboid-like) microglia is both necessary and sufficient to induce masculinization of the synaptic profile as a result of elevated PGE2 (*5-6*). We now report the surprising finding that mast cells, which are bone marrow-derived innate immune cells activated by peripheral allergy, anaphylaxis and atopy (*7*), are key players in brain sexual differentiation via communication with microglia.

## Results

We found that there are more mast cells in the POA neuropil and adjacent leptomeninges of neonatal male rats than females during the critical window for sexual differentiation (Fig. 1A-B). Thereafter, the mast cell population in males decreases to female levels (Fig. 1B). To determine if this sex difference is hormonally mediated, newborn females were treated with a dose of 17β-estradiol that mimics the levels normally seen in males (*8*) and the effect on mast cells was measured within 2 days. Mast cells are located in the neuropil and meninges of the POA as well as in the velum interpositum near the hippocampus (Fig. 1C). The overall number of mast cells (Fig. 1D) and the number in the neuropil (Fig. 1E) were increased to male levels in females treated with a masculinizing dose of estradiol. Mast cells release secretory vesicles filled with bioactive mediators via a process called degranulation (*7*). The number of mast cells with evidence of degranulation was not different between males and females on postnatal day (PN) 2, which is after the fetal male androgen surge, but exogenous estradiol induced acute increases in the percent of total mast cells showing degranulation in females (Fig. 1D, shaded bars). In agreement, males had a higher percentage of mast cells with evidence of degranulation than females on embryonic day 20, when estradiol is endogenously high in males due to the testicular hormone surge (Fig. 1F, shaded bars). There was no evidence of mast cell proliferation in the neonatal POA of either males or females (Fig. 1, S1A). Thus, sex differences in mast cell number appear to result from hormone-induced recruitment into the POA from nearby regions or the periphery as opposed to local proliferation. POA mast cells stain positive for all mast cell markers that were assessed, including histamine, serotonin, mast cell protease 2, the Immunoglobulin E (IgE) receptor, FCER-1, Alcian Blue and Safranin (Fig. 1, S1B), and in vivo counts of mast cell numbers using serotonin staining corroborate toluidine blue counts (Fig. 1, S1C). Estrogen receptor (alpha isoform) was detected in ~40% of mast cells in males and females (Fig. 1G-H). Estradiol treatment of isolated brain mast cells *in vitro* induced release of histamine (Fig. 1K) but had no effect on release of membrane derived PGE2 or prostaglandin D2 (PGD2) (Fig. 1I-J and Fig. 1, S2A-C), suggesting the effects of estradiol are specific to degranulation.

**Figure 1.**
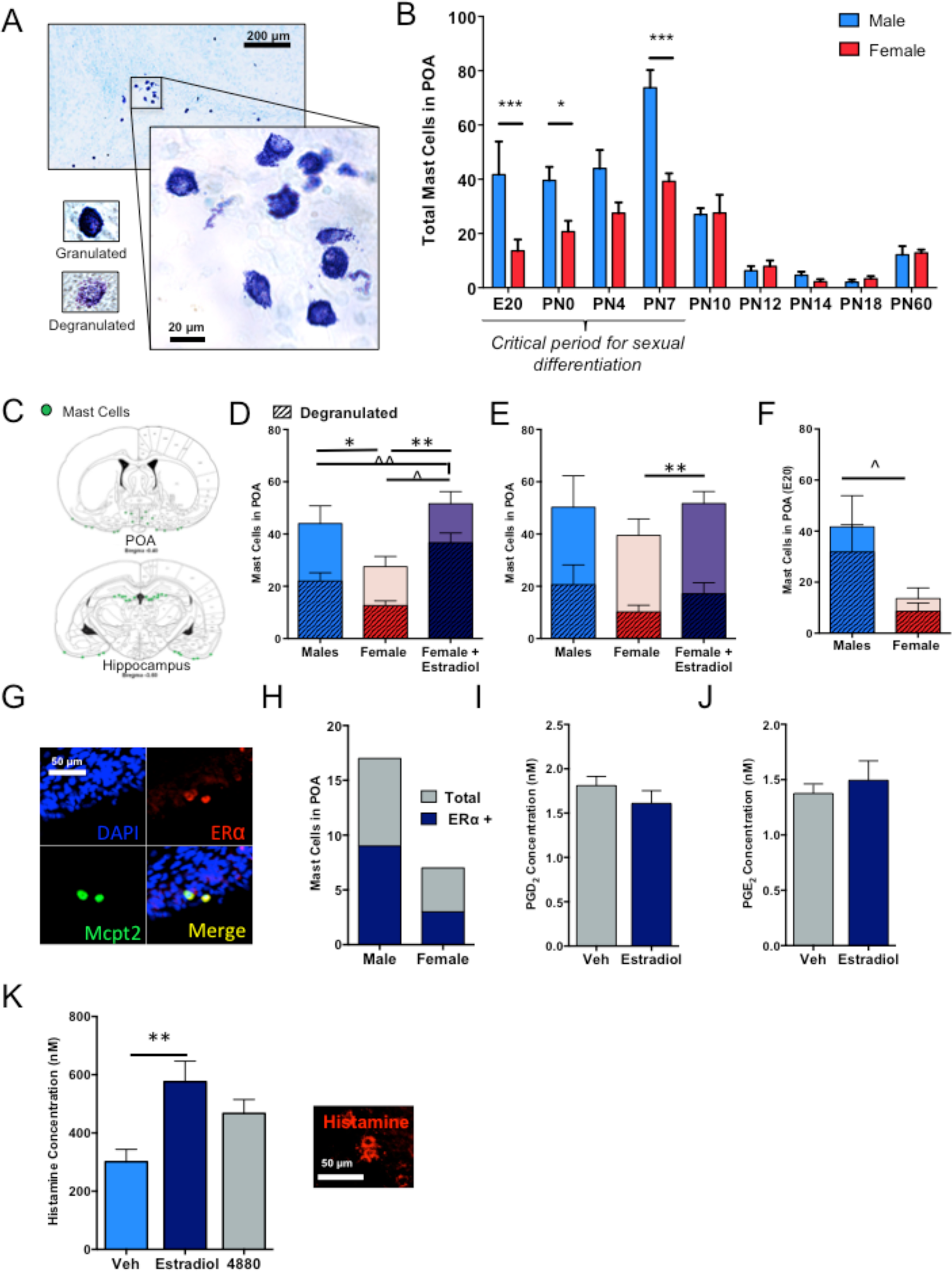
Sex differences in mast cells in the POA during the critical period for sexual differentiation. (A) Mast cells in the POA neuropil and leptomeninges on postnatal day 0 stained with toluidine blue. (B) Males had more mast cells than females on E20, PN0, and PN7, the critical period for sexual differentiation, [F_int(8, 106)_= 5.1, *p*<0.0001]. Mast cell numbers significantly increased until PN7 and decreased thereafter in both sexes with no further sex differences. (C). Schematic of mast cell location in the forebrain (green circles). Mast cells in the POA are situated in both the neuropil and leptomeninges, and additional mast cells are located in the velum interpositum adjacent to the hippocampus. (D-E) Treatment of females on PN0-1 with a masculinizing dose of estradiol increased (D) the total number of POA mast cells [F_(2,17)_=6.20, *p*=0.01; full bar graph] and the percentage of degranulated mast cells [F_(2,17)_=6.24, *p*=0.01; shaded portion of bar graph shows number of degranulated cells] (E) and the number of mast cells in the neuropil (shaded bars) versus the meninges [F_(2,17)_=9.22, p=0.002] on PN4. (F) On E20, during the testicular androgen surge in males, males had a higher percentage of degranulated mast cells than females [t_(9)_ = 2.51, p=0.03; shaded bars depict number of degranulated cells). (G-H) Approximately half of POA mast cells stained with mast cell protease 2 (Mcpt2) antiserum also stained positive for estrogen receptor alpha in males and females. (I & J) Estradiol treatment did not increase the amount of PGD2 (I) or PGE2 (J) released from cultured mast cells isolated from the postnatal brain. (K) Mast cells stain positive for histamine in vivo, and treatment of primary cultured mast cells with estradiol increased histamine release [F_(2,29)_=6.05, *p*=0.01]. Data presented as mean ± SEM. *indicates p<0.05, **indicates p<0.01, **** indicates p<0.0001 for full bar graphs; ^ indicates p<0.05 for percent total shaded bar graphs, ^ ^ indicates p<0.01 for percent total shaded bar graphs. Group sizes: B: E20: ♂ n= 6, ♀ n=5; PN0: ♂ n=8, ♀ n=7; PN4 ♂ n=6,♀ n=7; PN7: ♂ n=7, ♀=5; PN10: ♂ n=7, ♀=5; PN12: ♂ n=10, ♀ n=10; PN14: ♂ n=6, ♀=8; PN18: ♂ n=7, ♀=8; PN60: ♂ n=6, ♀ =6. D/E: ♂ n=6, ♀ n=7, E2-♀ n=7. D: ♂ n=10, ♀ n=10, E2-♀ n=7. F: ♂ n= 6, ♀ n=5. G/H: ♂ n=3, ♀ n=3; I/J: n=3 per time point. K: n=10 per group (two experiments with n = 5 for each).

Mast cells are frequent partners with microglia, the primary immunocompetent cells of the brain (*7, 9*). Microglia are emerging as important guides of developmental plasticity (*10*), including a role in establishing sex differences in the POA (*5*). Microglia in the neonatal male POA are more numerous, proliferate at a higher rate (Fig. 2, S1A-D), are more ameboid in shape, and produce more PGE2 than the highly ramified microglia of females (*5*). Therefore, assessing microglial morphology following mast cell manipulations is an excellent readout for downstream masculinization of the POA. In order to determine if mast cells are communicating with microglia to establish this sex difference, we used a rapid-acting pharmacological strategy to induce mast cell degranulation, by infusing Compound 48/80, a potent and validated mast cell secretagogue (*11-13*), directly into the brain of newborn females and subsequently assessed microglia morphology in the POA at 2 days of age. Compound 48/80 significantly increased mast cell degranulation in the POA (Fig. 2, S2A) and led to rapid increases in histamine turnover in the POA (Fig. 2, S2B). In primary POA cultures containing neurons, astrocytes and microglia but no mast cells, Compound 48/80 had no off-target effects directly on neuronal spine formation (Fig. 6, S2), indicating that effects of Compound 48/80 are mast cell dependent. We found a significant sex difference in the percent of ameboid shaped microglia in control animals, and stimulating mast cell degranulation in females resulted in a complete sex reversal of microglial morphology (Fig. 2A-B). Compound 48/80 did not alter mast cell degranulation in the spleen (Fig. 2, S2C), confirming effects were restricted to the CNS. To determine whether masculinization can not only be induced by mast cell degranulation in females, but also prevented by mast cell inhibition in males, we utilized a prenatal treatment regimen with the FDA-approved mast cell stabilizer, ketotifen, and assessed mast cell and microglia endpoints. In utero exposure to the mast cell stabilizer, ketotifen, via maternal drinking water decreased the percent of degranulated mast cells in the offspring POA (Fig. 2C) and also reduced the percent of ameboid shaped microglia in males (Fig. 2D). Microglia in the POA stain positive for histamine receptors type 1 and 4 (Fig. 2E), suggesting that mast cells, when stimulated to degranulate by estradiol or Compound 48/80 (Fig. 1K) may signal to microglia via histamine.

**Figure 2.**
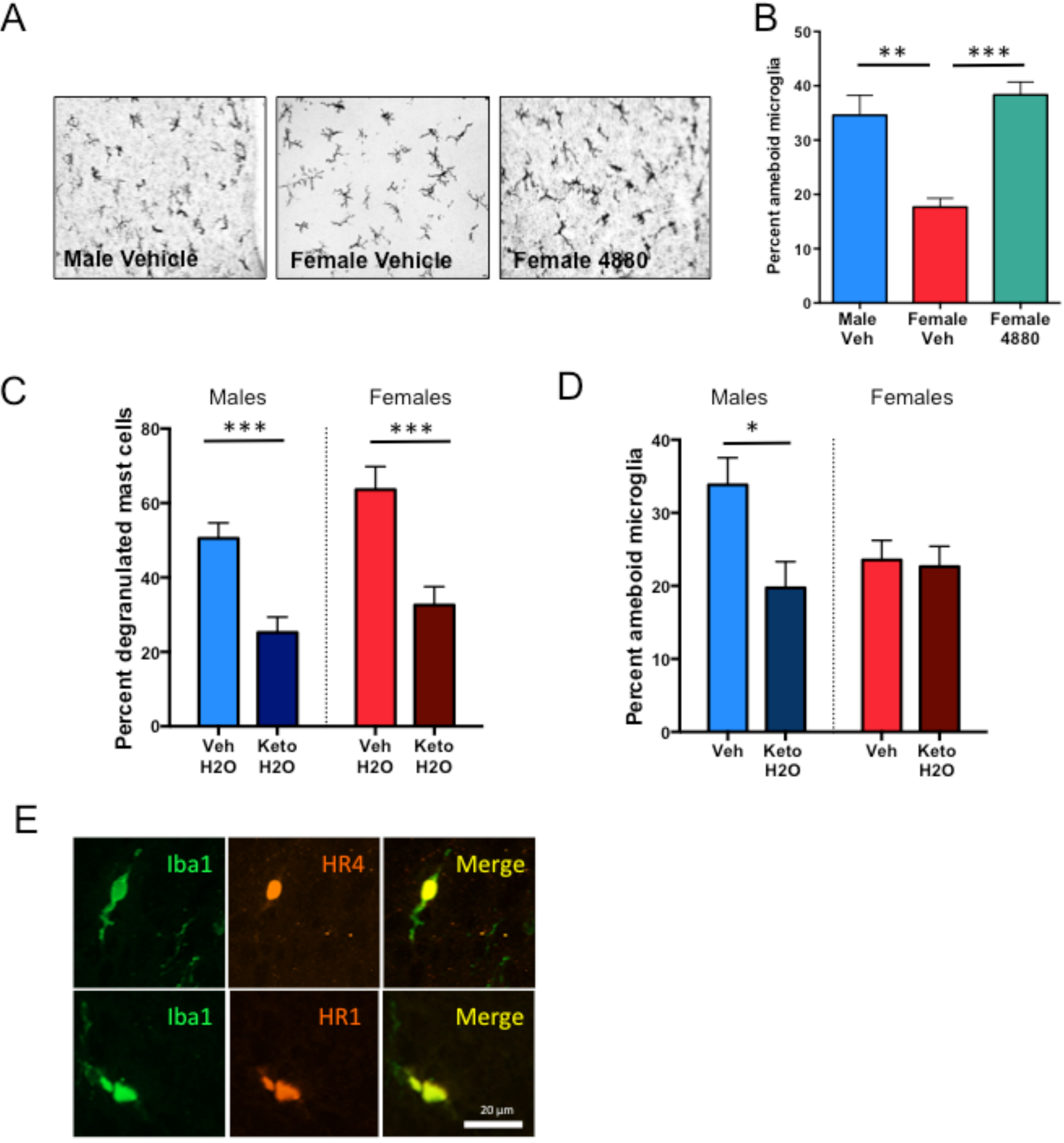
Effects of perinatal mast cell manipulations on innate immune cell number and activation in the neonatal POA. (A & B) Treatment of females with the mast cell degranulating agent Compound 48/80 increased the proportion of microglia in an ameboid state in females to male levels [F_(2,15)_=12.44, *p*=0.0007]. (C-D) Prenatal treatment of maternal dams with the mast cell stabilizer, ketotifen, in their drinking water led to significant decreases in (C) the percent of degranulated mast cells in the POA of offspring of both sexes [F_(3,19)_=32.62, *p*<0.0001; HSD p<0.01] and (D) decreased the proportion of microglia in the ameboid state in males to the level seen in females [F_(3,19)_=3.35, *p*=0.0.057; HSD p=0.038)]. (E). Iba1+ Microglia within the POA co-stain for histamine receptors type 4 (top panel) and type 1 (bottom panel). Data presented as mean ± SEM. *indicates p<0.05, **indicates p<0.01, **** indicates p<0.0001. Group sizes: Panel B: ♂V n=7; ♀V n=5; ♀4880 n=6. Panel C-D: ♂V n=5, ♂ Keto n=5, ♀V n=5; ♀ Keto n=6.

To determine whether a physiologically relevant immune challenge could similarly influence sexual differentiation of the POA, we developed an allergic challenge model that induces IgE production and activates mast cells. Adult female rats were sensitized to the allergen ovalbumin (OVA), bred, and challenged intranasally with OVA in saline or saline vehicle on gestational day (GD) 15 of pregnancy (Fig. 3A), which induced a significant acute IgE response in maternal dams (Fig. 3B). When offspring were assessed on PN4, control males again had more mast cells in the POA than control females, but gestational allergic challenge significantly increased the number of mast cells in the female POA to male-typical levels (Fig. 3C). Prenatal allergic challenge also increased the proportion of degranulated mast cells (Fig. 3D) and increased the percentage of microglia with ameboid shaped morphology in females to male-typical levels (Fig. 3E). There was a trend towards the opposite effect in males with a reduction in ameboid shaped microglia (Fig 3E), suggesting that at least on the short-term the response to in utero allergic challenge is opposite in males and females.

**Figure 3.**
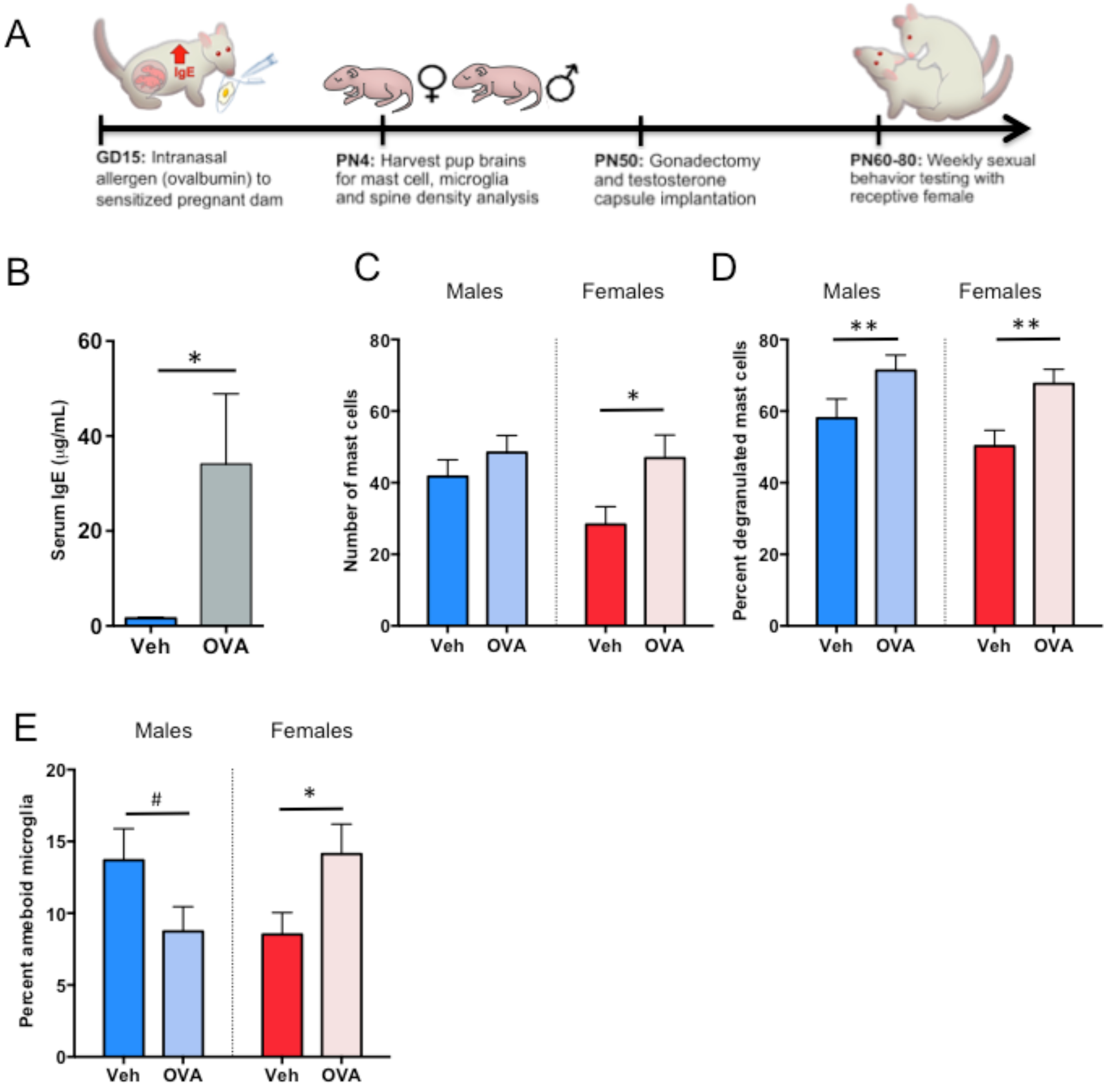
Effects of prenatal allergic challenge on mast cells and microglia in the POA. Ovalbumin (OVA) sensitized or control females were bred and challenged intranasally with OVA or vehicle on GD15 (A), and pups assessed neonatally for brain-resident immune cell numbers or grown to adulthood for sexual behavior testing (Fig. 3C-E; Fig. 4G-I). (B) Intranasal administration of OVA to pregnant dams at GD15 increased serum IgE levels in dams, confirming the generation of an allergic response [t_(4)_=2.20, *p*=0.046]. (C) Exposure to prenatal allergic challenge increased mast cell number in the POA of newborn females relative to controls, [F_(1,50)_=5.36; *p*=0.025]. (D) Allergic challenge also increased the percent of mast cells that were degranulated in both sexes, [F_(1,50)_=10.73, *p*=0.002], (E) but impacted microglia in males and females differently [F_int.(1,21)_=7.76, *p*=0.01]. Allergic challenge increased the percentage of ameboid microglia in the female POA relative to controls (HSD p=0.03) but had a trend toward a significant decrease in ameboid microglia in males (HSD p=0.09). Data presented as mean ± SEM. *indicates p<0.05, **indicates p<0.01, **** indicates p<0.0001. .# indicates trend 0.05≤p<0.1. Group sizes: Panel B: All groups n=3. Panel C-D: ♂V n= ♂OVA n=15, ♀V n=15, ♀OVA n=14. E: ♂V n=4, ♂ OVA n=7, ♀V n=8, ♀OVA n=6.

To determine whether the impact of greater numbers and more active mast cells on microglial morphology also influences early life programming of male-typical copulatory behavior, newborn female pups were treated with the mast cell stimulator, Compound 48/80, and male and female control pups were treated with saline vehicle on postnatal days 0-1. Pups were then grown to adulthood, gonadectomized, implanted with testosterone capsules to provide adult male-typical hormone levels and assessed for their motivation (latency) and execution (mount rate, ejaculation frequency) of masculine copulatory behavior with a sexually receptive female. The 1^st^ trial familiarized animals to the test paradigm while the 2^nd^ trial a week later was used for assessment of behavior. Treatment with Compound 48/80 masculinized the behavior of females, as they started mounting more quickly (Fig. 4A) and more often (Fig. 4B) compared to female controls, such that their overall performance was equivalent to males.

**Figure 4.**
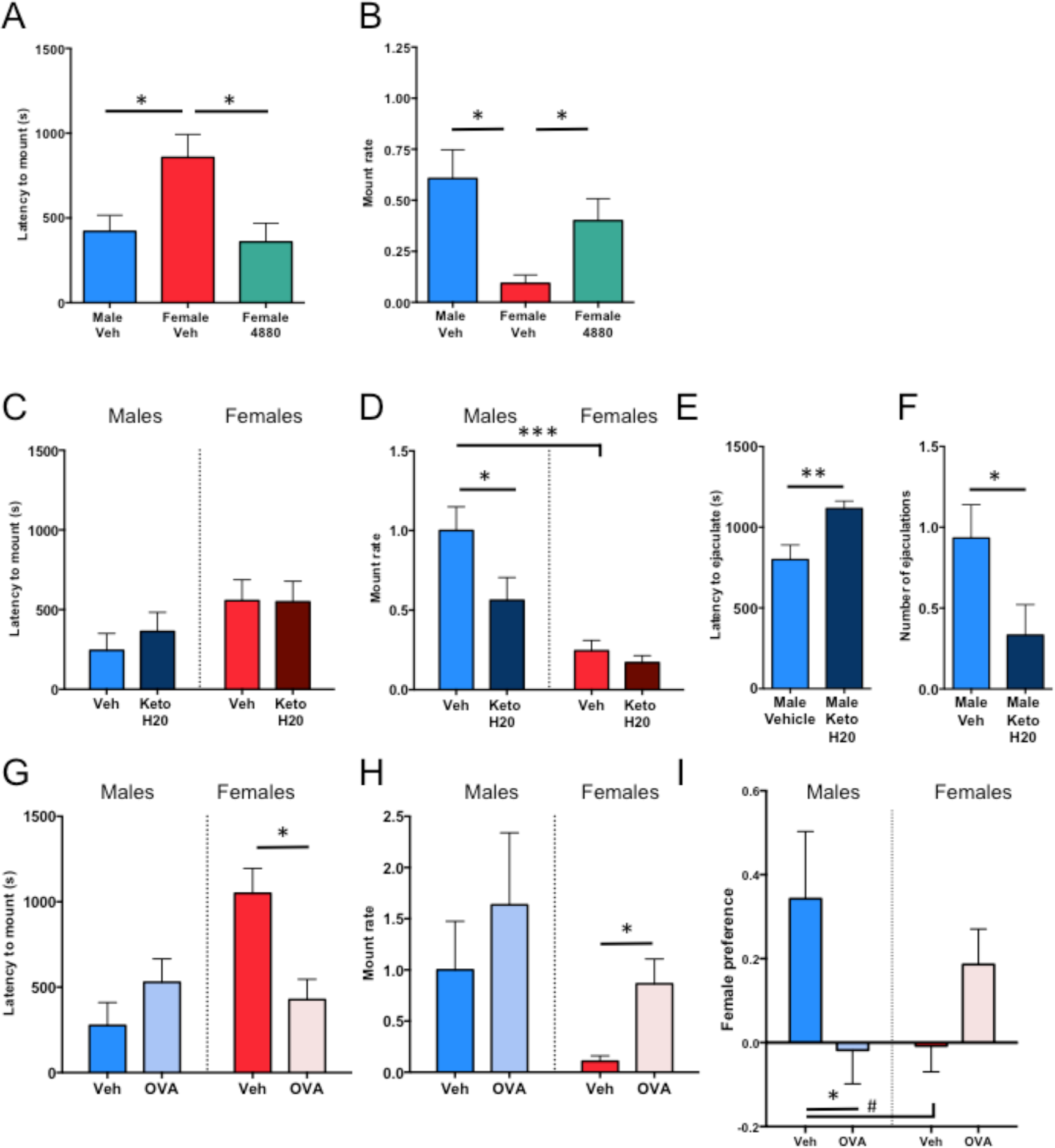
Manipulation of mast cells perinatally modifies male-typical reproductive behaviors in adulthood. (A-D) Adult females treated postnatally with compound 48/80 began mounting more quickly (A: F_(2,35)_=5.12, p=0.011, ♂V - ♀V HSD p=0.017; ♀V-♀4880 HSD: p=0.023) and mounted more frequently (B: M-W U=20, p=0.013) than control females. (C). Although adult males treated perinatally with ketotifen (given via the pregnant dam’s drinking water between GD17 and PN7) did not delay in mounting (C), they nonetheless mounted less frequently (D: M-W U=34, p=0.013), reached their first ejaculation later (E: M-W U=130.5, p=0.0135) and ejaculated fewer times (F: M-W U=37.5 p=0.021) than control males. (G-I). Adult females exposed to a maternal allergic challenge *in utero* began mounting more quickly (G: M-W U=45, p=0.013) and mounted more frequently (H, M-W U=5, p=0.013) than control females such that they were indistinguishable from control males. (I) Generally males prefer to investigate bedding soiled by rats of the opposite sex whereas females generally show no bias in preference, unless they are in estrus. Here, males trended toward a significant bias in odor preference compared to female counterparts [F_(1,23)_=21.81, p=0.016, HSD p=0.063]; however, odor preference was abolished in males experiencing allergic challenge *in utero* relative to unchallenged males (p=0.045). Allergic challenge *in utero* had no significant effect on odor preference of female littermates (p=0.2). Data presented as mean + SEM. # indicates trend 0.05≤p<0.1. *indicates p<0.05, **indicates p<0.01, ***indicates p<0.001. Group sizes: Panel A-C: ♂V n=17, ♀V n=10; ♀ 4880 n=11. Panel D-G: ♀ V & ♀Keto n=12, ♂V n=15, ♂Keto n=11. Panel H-J: ♀ Ova n=9, ♀ V n=5, ♂Ova n=9, ♂ V n=6.

We next sought to determine the converse of the above experiments that demonstrate that mast cell activation in females is sufficient to induce masculinization, specifically to show that mast cell inhibition prevents masculinization in males, which largely occurs prenatally. Pregnant dams were administered the mast cell stabilizing drug ketotifen in their drinking water. There was no effect of prenatal ketotifen exposure on the latency of adult male offspring to mount sexually receptive females (Fig. 4C) but the mount rate was significantly reduced in ketotifen-exposed males compared to controls (Fig. 4D). These same males took longer to ejaculate (Fig. 4E) and ejaculated less frequently (Fig. 4F) compared to non-ketotifen exposed males.

Using the prenatal allergic challenge model, we found that females exposed to a prenatal allergic challenge and then treated with testosterone in adulthood started mounting more quickly (Fig. 4G) and mounted more often (Fig 4H) than non-exposed females, reflecting a high degree of motivation and behavioral execution in the OVA allergen exposed females. Thus the same allergic challenge that suffices to recruit mast cell numbers and degranulation to male levels (Fig. 3) can also masculinize behavior. Females born to allergic challenged dams also trended towards a preference for olfactory investigation of female versus male soiled bedding (Fig. 4I), further indicating behavioral masculinization. Males gestated in an allergic challenged dam showed the opposite, a loss of female partner preference, suggesting dysmasculinization (Fig. 4I). Overall, these data are consistent with the view that mast cells are crucial regulators of early life programming of adult sex-specific behavioral repertoires, and suggest that both prenatal and postnatal effects of mast cells contribute to sex-specific behavioral programming. However, additional components of the allergic response may contribute to dysmasculinization in males and/or masculinization of females.

Neurons in the POA exhibit a marked sex difference with males having 2-3X the density of dendritic spine synapses and higher levels of the dendritic spine marker protein, spinophilin, compared to females (*5*). Both measures positively correlate with motivation for sex and frequency of male-typical mounting (*8*). Neonatal females gestated in an allergic-challenged dam had male-like dendritic spine density on Golgi-Cox impregnated POA neurons, while neonatal males from those same dams exhibited signs of dysmasculinization as density of dendritic spines was reduced (Fig. 5A-B). However, while the masculinization of dendritic spine density observed in neonatal females endured into adulthood (Fig 5C), the decrease observed in neonatal males was no longer evident in adulthood, with males gestated in allergic-challenged dams being indistinguishable from control males (Fig 5C). Further evidence that it is mast cells that direct the masculinization of neuronal morphology is found in the effect of postnatal treatment of newborn females with the mast cell secretagogue, Compound 48/80 which increased spinophilin protein levels in the POA to masculine levels (Fig. 5D; Fig. 5, S1C). There was no sex difference or effect of allergic challenge on overall dendritic length or neuronal cell body size (Fig. 5, S1A-B). In contrast to the masculinizing effects of prenatal allergen exposure on dendritic spines, postnatal immune challenge on the day of birth with lipopolysaccharide (LPS) did not lead to masculinization of spinophilin content in the POA on PN4 (Fig. 6, S1), indicating some specificity of allergic inflammation in particular on the masculinization process in the POA.

**Figure 5.**
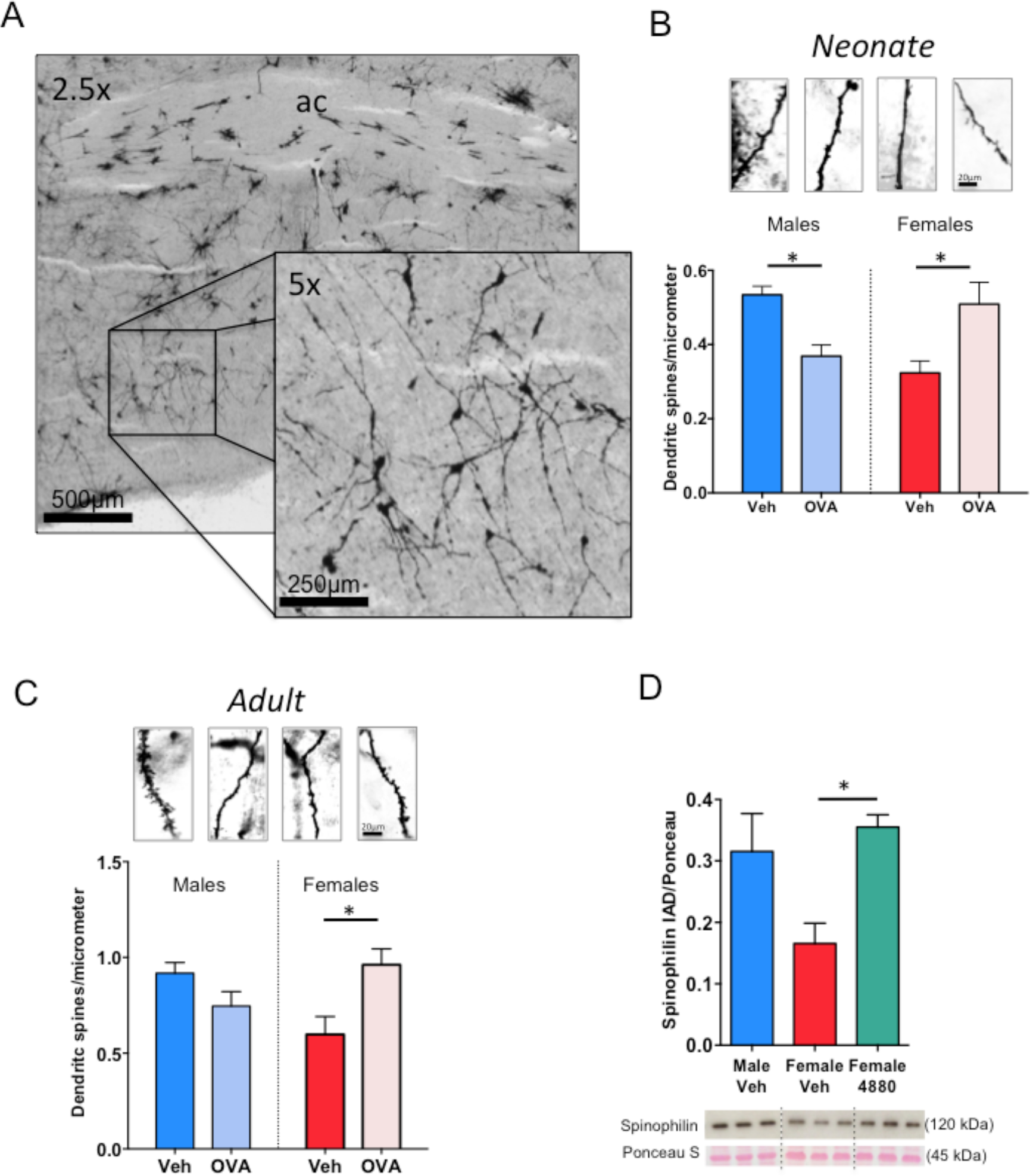
Prenatal allergic challenge and mast cell induced masculinization of POA dendritic spines and spinophilin in vivo. (A-B) Prenatal allergic challenge increased the density of dendritic spines in Golgi-Cox impregnated POA neurons of females on PN5, and decreased the density on male POA neurons [F_int[1,12)_=20.92, *p*=0.0006; Bonferonni post-hoc p’s < 0.05]. (C) Prenatal allergic challenge led to lifelong masculinization of dendritic spine density of females relative to vehicle females, as measured in adult POA neurons visualized with Golgi-Cox impregnation [F_int(1,12)_=11.24, p=0.006, HSD p=0.016], but had not enduring effect on males (D) Treatment with mast cell secretagogue compound 48/80 on PN0-1 increased spinophilin protein levels in the neonatal female POA on PN2 relative to control females, [F_(2,17)_=5.15, *p*=0.02]. Group sizes: B: All groups n=4. C: ♂V n=4; ♀ V n=3; ♂V n=4; ♀OVA n=5. D: ♂V n=7; ♀V n=6; ♀4880 n=7.

To establish if the effects of mast cells on neuronal dendritic spines are local to the POA and depend on microglia, we generated sex-specific primary POA mixed cultures and removed microglia from a subset and mast cells from all. Separately, isolated mast cell cultures were treated with Compound 48/80 and conditioned medium applied to the primary POA cultures. (Fig. 6D) Two days later spine-like protrusions on neurites were quantified. Conditioned medium from stimulated mast cells increased the density of spine-like protrusions on female POA neurons, but only if microglia were present in the cultures (Fig. 6A). Direct treatment of neuronal cultures with the Compound 48/80 had no effect, confirming the role of mast cells and indicating no non-specific effects on neurons or astrocytes (Fig. 6, S2).

**Figure 6.**
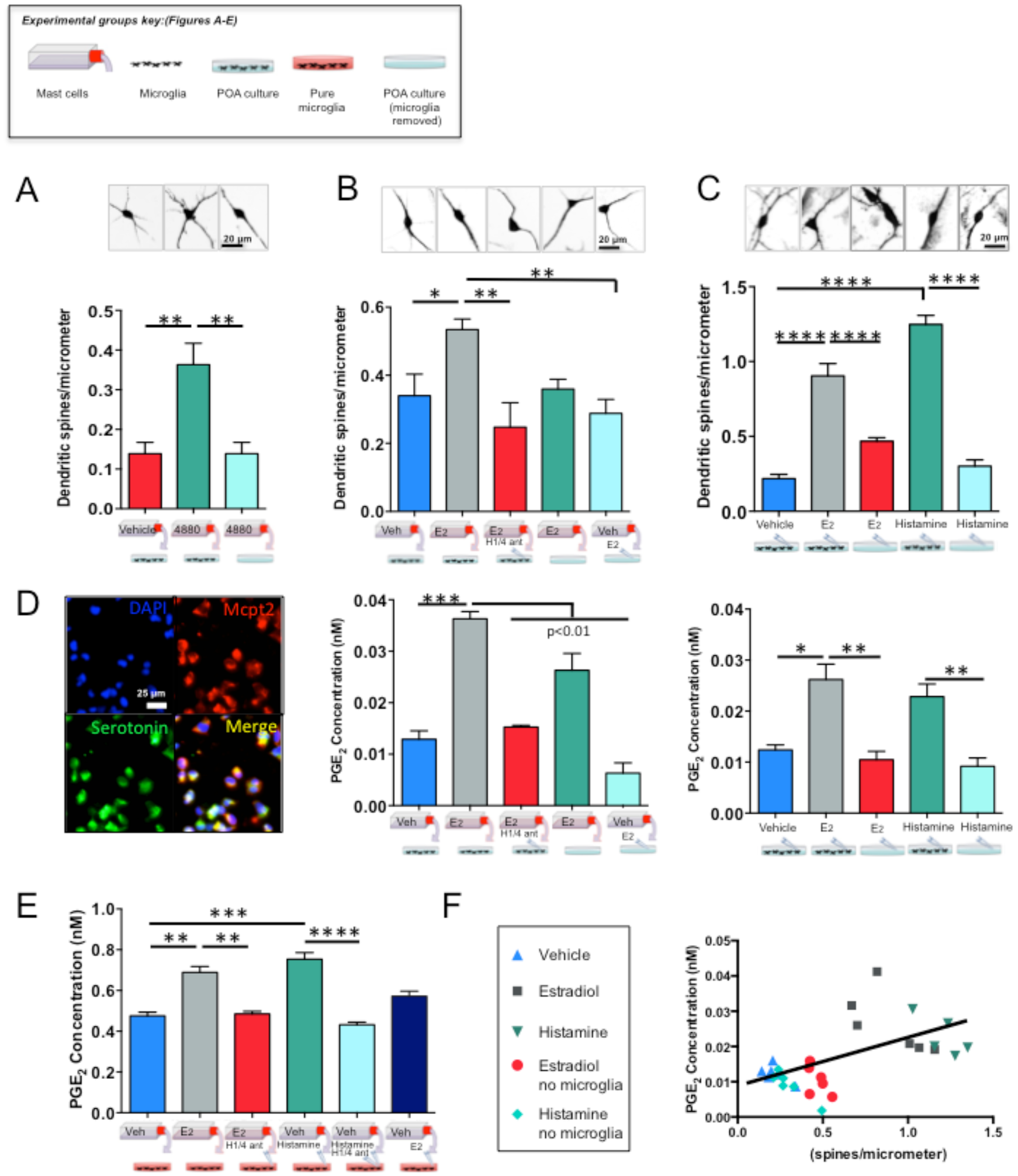
Mechanisms of mast cell induced masculinization of POA dendritic spines in vitro. (A) The density of dendritic spine-like protrusions on female primary POA neurons increased after exposure to conditioned medium from mast cells stimulated with compound 48/80, but prior removal of the microglia prevented this increase [F_(2,9)_=11.22, *p*=0.0036]. (B) Likewise, treatment with conditioned media from estradiol-stimulated mast cells increased the density of dendritic spines (top graph, [F_(4,29)_=7.717, *p*=0.001, HSD *p*=0.024]) and PGE2 concentration (bottom graph, F _(4,29)_=43.82, *p*<0.001, HSD *p*<0.001), and prior removal of microglia or inhibition of H1 & H4 histamine receptors prevented these increases (*p*=0.001). (C) Administration of histamine directly onto primary POA cultures also increased the density of dendritic spines [top graph, F_(4,29)_=66.09 p<0.001, Tukey’s HSD p<0.001 all] and PGE2 concentration [bottom graph, F_(4,29)_=12.43 p<0.001, Tukey’s HSD p<0.001 & p=0.027 respectively] to levels equivalent to that induced by estradiol treatment, and again prior removal of microglia prevented these increases. (D) Validation of mast cell pure primary cultures. All DAPI positive cells (top left) from primary mast cell cultures stain positive for mast cell protease-2 (Mcpt2; top right) and serotonin (bottom left), (bottom right; merged). (E) PGE2 concentration increased in microglia cultures after exposure to histamine or conditioned medium from mast cells stimulated with estradiol [F_(6,44)_=27.82 p<0.001, Tukey’s HSD p<0.001], but not following pretreatment with H1 & H4 histamine receptor antagonists. (F) Dendritic spine levels positively correlate with PGE2 concentration (R^2^=0.318, *p*=0.001) in primary POA cultures which were treated with vehicle, estradiol, histamine, or in which microglia were removed from the cultures before addition of estradiol or histamine. Data presented as mean + SEM. *indicates p<0.05, **indicates p<0.01. All photomicrographs and immunoblots are representative of treatment group. Group sizes: A: All groups n=4. B: All groups n=6. C/F: All groups n=6. E: V, MC-E2, MC-E2 + HA, & H n=8, H + HA n=7, HA & E2Direct n=6.

Given that mast cells release a spinogenic factor that is not PGE2 (Fig. 1I-J; Fig. 1, S2), estradiol increases histamine release from mast cells in vitro (Fig. 1K), microglia are necessary for propagating the mast cell derived signal (Fig. 6A) and microglia express histamine receptors (Fig. 2E), we hypothesized that histamine is the spinogenic factor released by mast cells in response to estradiol. Histamine then triggers microglia to produce PGE2, and PGE2 signals for spine formation (Fig. 1K). Consistent with this hypothesis, media from brain mast cells stimulated with estradiol increased spine-like protrusions on cultured POA neurons, but not if the primary cultures were pre-treated with antagonists for H1 and H4 type histamine receptors (Fig. 6B upper). Moreover, the media from mast cells stimulated with estradiol did not increase spine-like protrusions when microglia were removed from the primary cultures, and estradiol given directly to neuronal cultures devoid of microglia had no effect (Fig. 6B upper). Likewise, media from primary brain mast cells stimulated with estradiol triggered a two-fold increase in PGE2 levels in mixed microglia/neuronal cultures, but not if cultures were pretreated with H1 and H4 receptor antagonists or if the media was added to POA cultures in which the microglia were depleted (Fig. 6B lower), confirming mast cell-derived histamine acts on microglia to induce PGE2 which then induces masculinization of dendritic spine density on POA neurons.

Previously, we demonstrated that PGE2 induces spine-like processes on cultured POA neurons (4) and here we determined histamine does the same (Fig. 6C upper), by increasing PGE2 levels (Fig. 6C lower). If microglia were removed from POA cultures, histamine had no stimulatory effect on dendritic spine density (Fig. 6C upper) or PGE2 levels (Fig. 6C lower). The elevations in PGE2 content by histamine or estradiol-stimulated mast cell media could be recapitulated on isolated microglial cultures, but not if pretreated with H1 and H4 histamine receptor antagonists (Fig. 6E). Direct administration of estradiol onto POA cultures with microglia stimulated some PGE2 production but did not fully reestablish the elevations in PGE2 content observed in the mast-cell media that was stimulated with estradiol (Fig. 4G). While microglia are, thus, the primary source of PGE2 production in response to mast cell-derived histamine, there is also evidence of residual production of PGE2 by the neurons themselves (Fig. 6B lower), but it is nevertheless insufficient to significantly elevate spine formation (Fig. 6B upper). Levels of PGE2 and dendritic spine density were positively correlated across all groups in the *in vitro* histamine experiment (Fig. 6F), further confirming the spinogenic effect of PGE2. Moreover, treating newborn female pups with antagonists for histamine receptors types 1 and 4 reduced PGE2 production in the POA (Fig. 6, S2B), while treating newborn females with the mast cell degranulating agent, Compound 48/80, led to increased levels of spinophilin content in the POA, and co-treatment with H1 and H4 type histamine receptor antagonists prevented the masculinizing effects of mast cell activation (Fig. 7A). Overall, these data indicate estradiol acts in the developing POA to directly degranulate mast cells and stimulate the release of histamine; this mast cell-derived histamine activates microglia and spurs microglial production and release of PGE2; PGE2, in turn, promotes male-typical synaptic patterning on POA neurons which directs male-typical copulatory behavior in adulthood (Fig. 7B). The mechanism by which PGE2 stimulates dendritic spine synapse formation in the POA is known and involves EP2 & 4 receptors, activation of protein kinase A (PKA) and AMPA-type glutamate receptors (*14-15*). Excitatory afferent input to the POA from the medial amygdala integrates olfactory stimuli from sexually receptive females (*16*). Increased glutamatergic transmission in the POA is a prerequisite for male mating behavior (*17*). Thus the increased density of excitatory dendritic spine synapses on POA neurons in males assures sufficient excitation is achieved.

**Figure 7.**
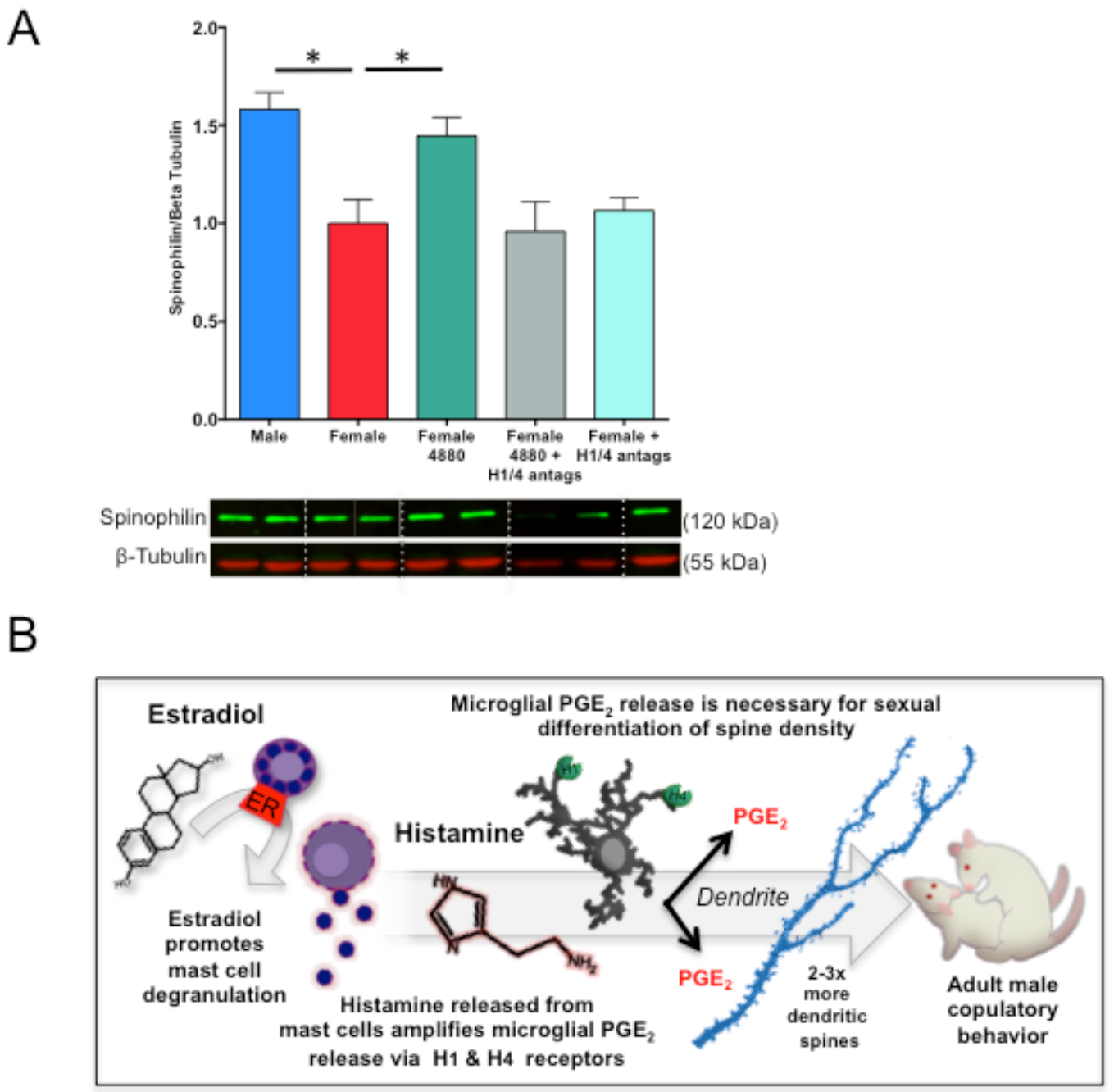
Effects of histamine receptor antagonism on dendritic spine proteins in vivo and working model of mast cell effects on the masculinization of the POA. (A) Treatment of neonatal females with Compound 48/80 (icv) led to increased spinophilin content in the POA, and co-treatment of females with histamine receptor 1 and 4 antagonists prevented this increase [F_(4,32)_=7.16 p<0.001, Tukey’s HSD p<0.001; Tukey’s HSD for M-F and F-F4880 p<0.05], indicating that histamine receptor activation is necessary for the masculinizing effects of mast cell degranulation on dendritic spines to occur in vivo. (B) Together, the current data suggest that estradiol acts on estrogen receptor alpha in mast cells to stimulate mast cell degranulation and the release of histamine into the male POA during the critical period for sexual differentiation. Histamine binds to H1 and H4 type histamine receptors on microglia and stimulates microglia to release prostaglandin E2 (PGE2). Microglia-derived PGE2 stimulates the masculinization of dendritic spine patterning on POA neurons in early life that enables male-typical copulatory behavior in adulthood. *indicates p<0.05. Group sizes: Panel A; ♂V n=9; ♀ n=8; ♀4880 n=7; ♀4880 +H1/4 antags: n=7; ♀H1/4 antags: n=6.

## Discussion

Steroid hormones are the principal drivers of brain sexual differentiation. There has been a long-standing assumption that hormones act largely upon neurons, as they express steroid receptors at high levels. We here demonstrate that sex-specific organization of the POA and resultant male-typical sexual behavior depends upon a surprising cellular player--the mast cell. Mast cells populate the brains of many species, including humans (*18*), but the possibility that mast cells are important to human brain development was discounted due to their sparseness. This warrants revisiting given our observation that a very small number of mast cells powerfully shape lifelong sex-specific endpoints in brain morphology and behavior.

Our data suggest that mast cells basally contribute to the masculinization process in males, as mast cell inhibition decreased male-typical sexual behavior in adulthood. Likewise, pharmacological or allergic inflammation-induced increases in mast cell activation in females can impact female sexual differentiation as well, shifting females toward a male-typical organization of the POA that is life-long and corresponds with increases in male-typical sociosexual behaviors. We further identified histamine as a mast cell-derived mediator that induces masculinization of microglia and dendritic spine endpoints, and likely impacts the masculinization process by stimulating microglia to release PGE2, which we have previously identified as the major molecular signal driving masculinization of the POA and male-typical sexual behavior (*4, 5, 15*). Depending upon their microenvironment, mast cells synthesize and release a host of mediators besides histamine, including serotonin, cytokines such as TNFα and interleukin 13, prostaglandin D2, and several proteases (*7*). These mediators may likewise contribute to the masculinization of the POA and warrant future inquiry.

Female brain development occurs in the absence of elevated levels of gonadal steroids and as a genetically programmed process should therefore occur along highly similar lines. Thus, girls would be predicted to show less variation in sex-typical behavior than boys, but the opposite is true (*19*). Likewise, girls with congenital adrenal hyperplasia experience prenatal androgen exposure and exhibit masculinization of some psychosexual traits (*20*), but the degree of behavioral masculinization does not track precisely with genital virilization, which is hormone dependent. We found females can be masculinized by an allergic response in the pregnant dam. While the impact of allergic response on the developing brain likely involves multiple signaling pathways and cell types beyond mast cells and their release of histamine, our results are consistent with an important role for mast cells as an additional and heretofore unknown source of variability in female behavior. A sex difference in the transcriptome of peripheral mast cells was recently identified (*21*), which if also true in the CNS could contribute to sex-specific impacts of prenatal allergic or inflammatory exposure.

Our findings highlight crosstalk between immune cells in the brain. Developing male rat brains have more microglia and mast cells and higher levels of inflammatory signaling molecules, such as PGE2. In humans, glial and inflammatory genes are more highly expressed in the fetal male brain (*22*), thus it is hypothesized that inflammatory mediators may serve the normal masculinization process in our species as well. Moreover, higher basal inflammatory signaling in the male brain may make males more vulnerable to altered behavioral development following inflammogenic perturbations early in life, including early life infection and stress (*23*). Indeed, our data indicates that males that experience allergic inflammation in utero show evidence of modest dysmasculinization in microglia, dendritic spines, and behavior. This may reflect an optimal threshold for inflammatory signaling in the male brain, beyond which there is dysregulated sociosexual development. Moreover, this dysmasculinization occurred even though allergen exposure further increased mast cell degranulation in males, which underscores that mast cells are likely just one of many potential modulatory systems that influence the sexual differentiation process in males.

A heightened basal innate immune cell presence in the male brain may also increase male vulnerability to neurodevelopmental disorders that count neonatal inflammation and infection amongst their risk factors, including autism, schizophrenia, and attention deficit hyperactivity disorder (*24-27*). Both autism and schizophrenia are associated with increased immune-associated gene expression and inflammatory signaling in the brain (*22, 24-27*). Autism is diagnosed four times more frequently in males (*28-29*) and schizophrenia varies in onset, symptomology and severity in men and women (*30-32*). The POA is rarely considered in the context of these disorders yet it is a crucial brain region for bonding, social recognition, and caregiving (*33-35*). Our findings uncover mast cells as a source of sex-specific variability in development that may contribute both to normal sex differences and to gender biases in risk for psychiatric and neurological disorders (*21, 35*).

## Materials and methods

### Animals

All experimental procedures were approved by the Institutional Animal Care and Use Committees at either the University of Maryland School of Medicine or The Ohio State University and conducted with Sprague Dawley rats purchased from Harlan Laboratories or bred in-house from animals originally from Harlan. Animals were housed on a reversed 12 h light/dark cycle in standard group cages, except when breeding, with food and water ad libitum. Adult females were paired with males and separated when vaginal lavage was sperm-positive. Once sperm-positive, pregnant females were isolated and allowed to deliver naturally. Rat pups used in experiments were birthed in-house, and pups from multiple litters were randomly assigned to experimental treatment and then randomly distributed back to dams to control for differences in maternal care. Cages were checked daily to determine the day of birth (designated postnatal day (PN) 0).

### In Vivo Treatments

#### Intracerebroventricular injections

Bilateral intracerebroventricular (icv) injections were performed under cryoanesthesia on PN0-1. A 23 gauge 1 μl Hamilton syringe attached to a stereotaxic manipulator was placed 1 mm caudal to Bregma and 1 mm lateral to the midline, lowered 3.0 mm into the brain, and then backed out 1 mm. One μl of drug or vehicle was infused over 60 s, and then the procedure was repeated on the other hemisphere. Compound 48/80 (Sigma; dose: 1 μg/ 2μl total) or 1μg/2μl total of a 50:50 mixture of 0.5μg/μl H1 and 0.5μg/μl H4 receptor antagonists (Centirizine and A943931 (Tocris)) was delivered in sterile saline vehicle and control animals were treated bilaterally with the same vehicle. The dose of Compound 48/80 used was determined based on previously published studies (*36*) and our own pilot studies showing its effectiveness at leading to mast cell degranulation (Fig. 2, S2A-B).

#### Subcutaneous and intraperitoneal injections

For other experiments, animals were treated subcutaneously with sesame oil vehicle or 17 *β*-estradiol (Sigma; dose: 100μg/ 0.1cc sesame oil) on PN0-1 and sacrificed on PN2. Bromodeoxyuridine (BrdU) is a thymidine analog incorporated into dividing cells for approximately 2 hr following injection. BrdU was administered intraperitoneally on PN1-2 (Sigma; dose: 100mg/kg in sterile saline) and animals sacrificed 6 hr following final injection on PN2. Lipopolysaccharide from E. coli (LPS; strain K-235, cat#L2143, Sigma; dose: 200μ/kg ip in 0.05 ml pyrogen-free saline) was given as an immune challenge on PN0, and control animals received an equivalent injection of saline vehicle. LPS-challenged animals were sacrificed on PN4. Tissue used in other experiments was collected at specified time points.

#### Gestational allergic challenge

Prior to pregnancy, adult females were sensitized with a subcutaneous injection of 1 mg ovalbumin (OVA grade V, Sigma) prepared at 4mg/ml in pyrogen-free 0.9% saline and precipitated at a 1:1 ratio with Al(OH)_3_ (Thermo Scientific) according to manufacturer’s instructions. After two weeks, a second 1 mg ovalbumin-Alum adjuvant injection was given. One week later, females were paired with males for breeding and detection of sperm assigned gestational day 0 (GD0). At GD15, pregnant rats were challenged intranasally with 1% ovalbumin in saline or saline vehicle (50μl per nare), which was placed on each nare under light isoflurane anesthesia and inhaled upon regaining consciousness. At 30 min following challenge, maternal blood was collected to assay for total serum Immunoglobulin E (IgE), using an IgE Rat ELISA kit (Abcam cat#157736) and serum samples run in triplicate. Females were paired in groups of two until GD15 and then housed individually. After birth, animals were sacrificed via perfusion to analyze mast cells, neuronal morphology, or microglia in the brain, or were weaned at PN22 into sex-specific groups of three containing both OVA challenged and vehicle exposed offspring for behavioral testing.

#### Ad Libitum administration of ketotifen in pregnant dams’ drinking water

Pregnant dams were treated with mast cell stabilizer, ketotifen fumarate (Sigma, cat# K2628), added to ad libitum drinking water from GD17, through delivery, until PN7. Delivery times did not vary because of ketotifen treatment. Ketotifen treated dams drank an average of 36.75 ± 7.304 mL per day indistinguishable from vehicle controls (41.65 ± 7.532). Dam body weights before and after delivery were also indistinguishable, as were pup body weights. Final ketotifen doses averaged 26.3 ± 1.680 mg/kg/day to the dam.

#### In vitro primary cell isolations

##### Microglia and Mast cell isolation

Brain mast cell and microglia isolations were performed on male and female pups (PN2-PN3) using methods published previously (*37-38*) and included a whole brain tissue homogenization under a 15 min 0.5% trypsin (Invitrogen) and 1% DNASE digestion. After suspension in a 37%, 50%, 70% Percoll gradient and centrifugation under 1200 g for 40 min, microglia were taken from the interphase between 50% and 70%. Mast cells were later isolated using Percoll separation alone, by taking cells from below the 70% phase. Plating densities and mast cell culturing condition suggestions were taken from (*38*). Mast cells were plated or replated at a density of 7.5 × 10^4^ cells per well for a 24-well plate, 3 × 10^5^ per well for a 6-well plate in 0.5 mL or 2 mL respectively of DMEM/F12 50:50 medium (Cellgro) containing 10% FBS (Fisher), 1% Pen-Strep-Amphotericin (Quality Biological), 1% L-Glutamine (Cellgro) 100 ng/ml stem cell factor (SCF) (Peprotech), and 30 ng/ml rat IL-3 (R&D Systems), grown overnight and re-plated to remove contaminating adherent cells. Thereafter, mast cells were replated every 4 days and fed with media described above but without Pen-Strep-Amphotericin. Mast cells became adherent by 1 week post-isolation. Mast cells were replated 24 hrs prior to an experiment with media lacking in IL-3 and SCF at a density of 2.0 × 10^5^ cells per well for a 24 well plate or 12 × 10^6^ per 75 cm^2^ flask (Corning) containing 0.5 mL or 24 mL of DMEM/F12 50:50 medium (Corning cat# 16-405-CV) containing 10% FBS (Fisher), colony-stimulating factor-1 (CSF, 5 ng/mL), glutamine (2 mM). Microglia were then split 1:2 every 4 days and used within 2 weeks of isolation. Mast cell identity and purity of cultures was verified via toluidine blue staining and FIHC against serotonin and MCP-2 described above; whereas, microglia were verified with FIHC against CD11b (1:5000) Abcam ab75476.

##### POA primary culture

On PN0, the POAs from male and female pups (n= 6 – 10) were microdissected, placed into sex-specific tubes containing 2 ml of HBSS, and digested with 0.25% trypsin (Invitrogen) and 1% DNASE for 15 min at 37°C or with papain and enzyme A per the manufacturer’s protocol for the Neuronal Dissociation kit (Miltenyi Biotech). Supernatant was removed, and cells washed and gently triterated with a Pasteur pipette in plating media consisting of DMEM/F12 medium without phenol red, 5% FBS, and 1% antibiotic/antimycotic (all Invitrogen) until dissociated. Cell density and viability were determined on a hemacytometer using Trypan blue, and plated at 500,000 cells per 25 mm round poly-lysine coated coverslip in a 100μl volume in 30 mm round petri dishes. After allowing cells to seed for 2 hr, cultures were fed with 2 ml of cell culture medium, consisting of Neurobasal A Medium without phenol red (Invitrogen), 1% B27 supplement (Invitrogen), and 0.125% L-glutamine (Sigma), and allowed to acclimate and grow for 24 h before treatment. In some experiments, vehicle of 2-hydroxypropyl-B-cyclodextrin (HPCD) (Sigma, H107) or 10 nM estradiol in HCPD (Sigma, E4389) were added to all the cultures on DIV1. Histamine (1 μg/mL) or saline vehicle were also added to all the cultures. The same pharmacological treatments were repeated on DIV2 and 6 hr later, media was removed for PGE2 assay and replaced with 4% PFA for immunohistochemistry for Microtubule-Associated Protein-2 (MAP-2; Sigma cat# M1406, 1:1000, 24 hr), as detailed below, to detect spine-like protrusions from the dendrites. MAP-2 protein is not localized in dendritic spines but allows for accurate quantification of spine-like protrusions attributable to diffusion of the DAB reaction product from the neurites into the spine-like protrusion in cultured neurons (*15*).

##### Microglia-free POA cultures

POA primary cultures were depleted of microglia via agitation on the day after culture, day in vitro 1 (DIV1) via our previously published procedure (*14*), which depletes ~90% of microgila from cultures. Coverslips were shaken at 200 RPM for 30 min to detach microglia, after which the microglia-containing medium was replaced with fresh culture medium, and cultures treated as appropriate for the experiment. At the conclusion of the experiment, cells were fixed with 4% paraformaldehyde in PBS for 10 min, and processed for DAB-immunohistochemistry against MAP-2, as outlined below.

##### Mast Cell Conditioned Media Exchanges

Mast cells (8 × 10^6^) were plated in 24 mL in 75 cm^2^ flasks and IL-3 and SCF were removed 24 hrs prior to treatment. HPCD vehicle or 10 nM estradiol in HCPD were added for 30 min before addition of this media to primary mixed neuronal or isolated microglial cultures. To prepare the DIV1 sex-specific neuronal cultures, some of the cultures had microglia removed by shaking. Then some of the DIV-1 POA neuronal cultures or isolated microglia received 10μM cetirizine HCl (H1 receptor antagonist) and 10μM A94391 (H4 receptor antagonist) (Tocris, Bioscience, Cat#s 2577 & 3753, respectively) or saline vehicle controls. One-half or 1 mL of media was removed from the primary neuronal or isolated microglial cultures and replaced with 1 mL of treated mast cell media that was centrifuged at 300 g for 5 min to remove residual mast cells, and supplemented with either H1 & H4 receptor antagonists, estradiol, or vehicle depending upon the condition to ensure final concentrations remained 10μM and 10 nM respectively. The same treatments were repeated on DIV2 and 6 hrs later media was removed for PGE2 assay. Primary POA neuronal cultures were fixed with 4% PFA for immunohistochemistry for MAP-2 to detect spine-like protrusions from the dendrites.

### Analysis

#### Biochemical Measurements of PGE2 and PGD2

Mast cell, microglia, primary neuronal culture media or freshly dissected tissue from the POA were stored at -80° C for PGE2 (Arbor Assays, Cat# K051) and PGD2 (Cayman Chemicals, Cat# 500151) EIA. Prior to PGE2 assay, Tissue was homogenized in RIPA buffer containing 1% Igepal CA630, 0.25% deoxycholic acid, 1 mM EDTA, 154 mM NaCl, and 65 mM Trizma Base, with added protease and phosphatase inhibitors (1:1000). Protein supernatant was extracted after 20 minutes of centrifugation at 3000 rpm at 4°C, and total protein concentration determined via Bradford assay (BioRad). POA tissue samples were prepared for PGE2 extraction by acidifying 200uL of sample with 2N HCl to pH of 3.5. Samples were then placed on wet ice for 15 minutes and centrifuged for 2 minutes at 3000rpm to remove precipitate. C18 reverse phase columns (Thermo Scientific 60108-303) were prepared by washing with 10mL of 100% ethanol and then 10mL of dH2O. Isolute SPE Adapters (Biotage 120-1101) were used to fit the syringes onto the columns. PGE2 samples were run through the column and were then washed with 10mL of dH2O, 10mL of 15% ethanol, and 10mL of hexane. 2mL of Ethyl Acetate (Sigma) were used to elute the samples into a 12mL polypropylene tube. Tubes were placed in a rack at an angle to maximize surface area for evaporation of ethyl acetate. Samples were left in the fume hood for 48-72 hours to reach complete evaporation. All chemicals were obtained from Sigma unless otherwise specified. For PGE2 from neuronal cultures, the manufacturer’s high-sensitivity protocol was used and included loading 100 μL of sample in triplicate, whereas for tissue-extracted PGE2, microglial or mast cell media 50 μL of sample were loaded per the manufacturer’s standard protocol. Thereafter, 25 μL of PGE2-conjugate and 25 μL of primary antibody were used for a 2 hour incubation at RT followed by 4 rinses with wash buffer, 30 min of 100 μL TMB substrate development, and 50 μL 1 N HCl stop. Absorbance was read at 450 nm by a Tecan M1000 Infinite (Männedorf, Switzerland), and manufacturer standards regressed with a Marquat logistic-4 parameter fit. Extracted samples were corrected for total protein concentration and expressed as pictograms of PGE2 per milligram of total protein. For the PGD2 assay, 100 μL of cell culture media or manufacturers standards was combined with the 100 μL Methyl Oximating Reagent and 50 μL of this solution was added to each well in triplicate before addition of 50 μL of PGD2-MOX AChE tracer and 50 μL of primary antiserum and incubation for 1 hr at rt. After 5 rinses with wash buffer, 200 μL of Ellman’s reagent was added to each well and incubated for 60 minutes before reading absorbance at 410 nm and confirming a minimum 0.3 A.U. absorbance on the B_0_ well before manufacturer’s standards were regressed by Marquat logistic-4 parameter fit of the %B/B_0_.

#### Histamine release in vitro

Mast cells were plated in a 24-well plate at a density of 1.5 × 10^6^ cells/ mL and IL-3 and SCF was removed from the media 24 hr prior to treatment with a vehicle containing 2-hydroxylpropyl-B-cyclodextrin carrier (Sigma, Cat# H107) or 10 nM estradiol immersed in 2-hydroxylpropyl-B-cyclodextrin (Sigma, Cat# E4389). Thirty min. later, media was collected from each well, centrifuged at 300 g for 5 min to remove residual cells, and supernatant diluted at 1:50 prior to assessing histamine levels via EIA (Cayman Chemicals, Cat# A05890) per manufacturer’s protocol.

#### Histamine release in vivo

Standard 100 μM stock of Histidine, histamine, methylhistamine, and uniformly labeled ^13^C-glycine were prepared in ultrapure water. Brain samples were dissociated using sonication with tungsten beads, centrifuged, and amines extracted using boiling water (4 min incubation in hot water bath) followed by cold Acetonitrile/0.2% w/v Formic acid solution centrifugation, and evaporation of solvents using a speed vacuum. Dried samples were resusbended in 0.1% formic acid and analyzed using liquid chromatography tandem mass spectrometry (LC-MS/MS). The analysis of histamine, methylhistamine, histidine and 13C-Glycine (internal standard) was carried out using a UHPLC (Ultra High Pressure Liquid Chromatography) 1290 from Agilent Technologies, Inc. coupled to a hybrid Triple Quadrupole/Ion trap mass spectrometer QTRAP 5500 from AB Sciex.

Each pure standard was diluted to 1 *μ* M with a solution of acetonitrile/ultrapure water (50:50; v/v) containing 0.1% of formic acid. Metabolites were injected individually and directly into the mass spectrometer apparatus at a flow rate of 7 *μ* L/min. This step (e.g., direct infusion) optimizes four major parameters of a specific metabolite: i) the declustering potential (DP), ii) the entrance potential (EP), iii) the collision energy potential (CE), and iv) the collision cell exit potential (CXP).

The compound parameters were as follows:

| Metabolite | Parent Mass | Daughter Mass | Retention Time (min) | DP (volts) | EP (volts) | CE (volts) | CXP (volts) |
| --- | --- | --- | --- | --- | --- | --- | --- |
| Histamine | 112 | 95 | 3.43 | 70 | 10 | 19 | 14 |
| Histidine | 156 | 110 | 2.43 | 50 | 10 | 21 | 16 |
| Methylhistamine | 126 | 109 | 3.45 | 50 | 10 | 27 | 18 |
| $^{13}\text{C}$-glycine | 78 | 31 | 0.84 | 20 | 10 | 19 | 12 |

The extracts were transferred to glass vials and placed in an autosampler, kept at 4 °C. 20 *μ* L of sample were injected into the column. The liquid chromatography analysis was carried out at 30°C. Metabolites were resolved using a C18 Kinetex biphenyl (100 × 2.1 mm, 2.6 *μ* m) column and a security guard column (10 × 2.1 mm) from Phenomenex. The gradient used to separate the molecules consisted of ultrapure water with 0.1% formic acid plus 1 mM perfluoroheptanoic acid (solvent B) and 0.1 % formic acid in 100 % acetonitrile (solvent A). The total LC-MS/MS run was 8.5 minutes with a flow rate of 400 *μ* L/min. The gradient went as followed (solvent A): 0-1.0 minutes 10%, 1.0-4.0 minutes 50%, 4.0-4.1 minutes 90%, 4.1-6.0 minutes 90%, 6.0-6.1 minutes 10%, and 6.1-8.5 minutes 10%.

The mass spectra were acquired using Turbo Spray Ionization of 2500 V in positive ion mode and multiple reaction monitoring (MRM). The curtain gas (nitrogen), CAD (Collision Activated Dissociation), nebulizing and heating gas were set to 35 psi, medium, 40 psi and 40 psi, respectively. The temperature of the source was fixed at 650°C. The mass spectrometer was set to have a dwell time of 30 msec. LC-MS/MS data were acquired and processed using Analyst 1.6.1 software.

#### Histology and immunohistochemistry

For all *in vivo* histology experiments, animals were killed via lethal overdose with FatalPlus (Vortech Pharma) followed by transcardial perfusion with saline followed by 4% paraformaldehyde, brains removed, and postfixed for 12 hours. Brains were sectioned coronally at 45 μm thickness on a cryostat (Leica) and mounted onto SuperFrost charged slides (Fisherbrand) for subsequent staining procedures.

##### Mast cell staining

Mast cells were visualized using staining with acidic Toluidine Blue (0.5% in 60% ethanol; pH=2.0) as detailed in (*30*), sequential acidic Alcian Blue (1% in 0.7N HCl) and Safranin O (1% in 0.125N HC; all stains from Sigma), or immunohistochemistry for mast cell specific markers as detailed below.

##### Immunohistochemistry (IHC) and immunofluorescence (IF)

Brain sections were rinsed twice with PBS, permeabilized with 0.3% H_2_O_2_ in 50% methanol, blocked with 5-10% bovine serum albumin or normal goat serum in PBS + 0.4% Triton X, and incubated with primary antisera for 24-120 hours at 4°C. For *in vitro* experiments, coverslips were washed under PBS twice, and incubated with primary antisera for 24 h at 4°C containing 10% bovine serum albumin in PBS + 0.4% Triton-X. Thereafter, sections or coverslips were extensively washed, and either incubated with appropriate fluorescently-tagged secondary antibodies for 2 hr (AlexaFluors 488, 555, 594, or 633, Life Technologies; 1:250) for IF, or processed with biotinylated secondary antibodies (Vector), avidin-biotin complex (Vector), and reacted with diaminobenzidine with or without Nickel (DAB or Ni-DAB; Sigma) in 0.125M sodium acetate to visualize chromogen for IHC. IF tissue was subsequently incubated in Hoescht nuclear stain for 20 minutes (Invitrogen Cat#MP-10338; 1:5000). Stained sections were coverslipped with DPX mounting media or VectaShield Hard Set (Vector). Primary antisera used were as follows: Microtubule-Associated Protein-2 (MAP-2) (Sigma 1:1000, 24 h), Estrogen receptor alpha (ER-*α*) (Millipore Cat# 06-935, 1:10000, 120 h); Serotonin (Abcam Cat#sc32292, 1:1000, 24 h); Histamine (Abcam Cat#ab43870, 1:1000, 24 h); Iba1 (Wako Cat#019-19741, 1:1000, 24 h or Novus Biologics Cat#NB100-1028, 1:1000, 24 h); CD11b (Abcam Cat# ab75476, 1:5000, 24 h); The IgE receptor FC E R1 (Abcam Cat# ab33568; 1:1000, 24h); rat mast cell protease 2 (RMCPII) (Moredun Scientific Cat# MS-RM4, 1:10,000, 24 h) BrdU (BD Biosciences Cat#347580, 1:250, 24 h), histamine receptor 1 (EMD Millipore Cat# AB5652P; 1:5000, 24h), or histamine receptor 4 (Abcam Cat# ab97997; 1:5000, 24h). Microglia were quantified using Ni-DAB IHC, and proliferating microglia quantified using sequential DAB IHC against Iba1 and Ni-DAB against BrdU, and co-localization confirmed in three dimensions using confocal microscopy on IF-stained tissue (Fig. S2B-D). Mast cell phenotype, mast cell estrogen receptors, and proliferating mast cells were assessed using IF and imaged with confocal microscopy. For co-localization, primary ER-*α* and RMCPII antibodies were labeled with secondary antibody conjugated to AlexaFluor-594 and AlexaFluor-488 respectively.

#### Golgi-Cox staining

Whole brains from PN5 pups or P80-90 adults were placed in 15 ml of Golgi-Cox solutions A and B (FD Neurotech) for 10 days, then solution C (sucrose; FD Neurotech) for 1-1.5 weeks, cut into coronal sections 100 μm thick using a Leica vibrotome, and staining embedded using solutions D+E following the FD Neurotech protocol. Tissue was cleared with ascending ethanol, defatted with xylenes, and coverslipped using Permount.

#### Microscopy and Stereology

##### Stereology and single cell reconstruction

Either a Nikon Eclipse E600 Microscope or a Zeiss Axioimager.M2 microscope coupled to a CX9000 Digital Camera and Stereo Investigator software (MBF Bioscience) were used to estimate the total population of mast cells and microglia in the POA, using an average of 4-6 sections per animal encompassing the entire rostrocaudal extent of the POA. At the time of counting, mast cells were categorized as either granulated or degranulated, and microglia were categorized as ameboid or not ameboid. Microglia were considered ameboid if they had an enlarged cell body and either no processes or few, short processes, based on criteria validated in our previously published study (*5*). Neurolucida software (MBF Bioscience) was used for cell culture image acquisition and analysis. For MAP2-labeled cells and single-cell reconstruction, cells were reconstructed using a 100X oil objective and were chosen for analysis if they had at least 2 distinct processes and uniform dark labeling throughout the extent of the cell. Data collected included cell body size, total neurite length, number of neurites, and number of spine-like protrusions from the neurite. Labeled protrusions of < 5 μm in length from the neurite were counted as spine-like protrusions. Three-six cells per coverslip were used for the analysis.

For analysis of Golgi-Cox impregnated POA neurons, neurons were chosen for analysis if multiple processes were visible and the cell was easily distinguishable from nearby cells. 4-5 cells per animal across multiple brain sections were reconstructed in three dimensions under a 100x oil objective using Neurolucida software. Morphological parameters for each cell were computed using Neurolucida Explorer, including cell body size, total dendritic length per cell, number of dendritic segments and branch points, and total number of dendritic spines per neuron. For adult analyses, dendritic spines were analyzed, but single cells were not reconstructed in 3D. Dendritic segments chosen for analysis were unobstructed by other Golgi-stained material, and were at least 25 *μ* m in length without a bifurcation. Only one segment was analyzed per given cell. 4-5 dendritic segments per animal across multiple brain sections were reconstructed, and dendritic spine density was analyzed using Neurolucida Explorer software.

##### Confocal imaging

Double or triple labeled IF tissue was imaged using a Zeiss LSM 710 with a 20x Plan Apocromat 1.0 numerical aperture dipping objective and z-stacks acquired using Zen software to determine co-localization of Iba1 and BrdU staining (proliferating microglia) and serotonin and BrdU staining (proliferating mast cells).

#### Western blot

Tissue was homogenized in RIPA buffer containing 1% Igepal CA630, 0.25% deoxycholic acid, 1 mM EDTA, 154 mM NaCl, and 65 mM Trizma Base, with added protease and phosphatase inhibitors (1:1000). All chemicals were obtained from Sigma unless otherwise specified. Protein supernatant was extracted after 20 minutes of centrifugation at 3000 rpm at 4°C, and total protein concentration determined via Bradford assay (BioRad). Fifteen μg protein was electrophoresed on an 8-16% precast SDS polyacrylamide gel (Life Technologies) and transferred onto a single polyvinyl difluoride membrane (Bio-Rad). Membranes were blocked in 50% Odyssey blocking buffer (LI-COR) in TBS or 10% nonfat milk in 0.1% Tween in Tris-buffered saline (TTBS) and subsequently incubated with spinophilin primary antiserum (Millipore Cat#06-842, 1:1000) and Beta Tubulin primary anti-serum (Biolegend Cat# 801202, 1:3000) in 50% odyssey blocking buffer in TBST or in 5% milk in TTBS overnight at 4°C. Membranes were rinsed and incubated with IRDye anti-rabbit 800 and anti-mouse 680 produced in goat (1:20,000, LI-COR) in 50% odyssey blocking buffer in TBST containing 0.02% SDS in the dark or HRP-conjugated secondary antibody (1:200) for two hours. An Odyssey CLx imaging system was used for detection of fluorescent immunoblots. Spinophillin protein detected was normalized by the 55kDa beta tubulin band from each lane as a loading control. A Phototype chemilluminescence system (New England Biolabs) was used to detect the immunoblots by exposing the membrane to Hyperfield ECL (GE Healthcare). Integrative grayscale pixel area densitometry captured with a CCD camera was quantified with NIH Image software. Ponceau S staining appearing at 45 kDa was used as a loading control, and immunoblot densitometry values for each lane expressed as a percentage of Ponceau staining for the same lane.]

#### Behavior

Between PN50-54, animals were gonadectomized under isoflurane anesthesia and implanted subcutaneously with a 30-mm silastic capsule (1.57mm inner diameter, 3.18 mm outer diameter) filled with crystalline testosterone (Sigma) placed between the scapula. This capsule length mimics physiological levels of testosterone circulating in adult males and allows appropriate activational hormones for developmentally-masculinized females to perform male-typical copulatory behavior (*4*). Two weeks following surgery, animals were video recorded for at least 20 min during the dark phase of the light cycle, in a Plexiglass behavioral arena in the presence of a hormonally-primed receptive stimulus female under red light illumination. Behavioral data was quantified for every animal by an observer blind to the experimental treatment of each animal. Measures included number of mounts, latency to mount, ejaculation, time of each ejaculation and time to start mounting after each ejaculation. In order to record the full post-ejaculatory interval for situations during which a male started a post-ejaculatory interval with some of the 20 min left, the male was observed beyond the 20 min period until he started mounting again. These mounts outside of the 20 min period were not tallied in total mount measures. Mount rate could be calculated from the total mounts divided by the 20 minutes less the total time during that 20 min the animal was in a refractory post-ejaculatory state.

##### Olfactory preference test

Olfactory preference test was performed on adult animals (~PN60) using a modified protocol based on (*39*). Animals were placed in the same Plexiglass testing arena used for sexual behavior testing, with two ceramic dishes in opposite corners of the arena (counterbalanced across groups and animals). Dishes contained soiled bedding from adult male or female non-littermate conspecifics’ cages, freshly collected 5 days after bedding change. Animals were placed in the center of the arena and allowed to openly explore the arena for 5 min under red light illumination during the dark phase of the light cycle, while being videotaped. The number of seconds spent actively investigating (e.g., sniffing, digging in, or climbing on/in) each dish of soiled bedding was tallied and a female bedding preference score tabulated ([Time spent investigating female bedding- time spent investigating male bedding]/total investigation time).

### Data analysis

All histological, immunohistochemical, western blot, and culture data were analyzed using one- or two-tailed t-tests or one or two-way ANOVA, followed by Tukey’s HSD post-hoc tests, exclusions below. When normally distributed, behavior data were analyzed with one-way ANOVAs followed by Tukey’s HSD post-hoc analysis. When not normally distributed, behavior data were analyzed by Mann-Whitney U tests correcting *α* -significance for family-wise error. Chi-squared analysis was used for immunohistochemistry for co-localization of ER-alpha and mast cell protease 2. Overall statistical significance was set at *α* = 0.05. Exact p-values are presented in figure captions for each dataset and analysis. All data was analyzed using GraphPad Prism or SPSS software, and all graphs made using GraphPad Prism. Statistical results and group sizes are presented in each figure caption. Additionally, a summary of statistical details are listed in Table 1, including data structure, type of test, test value, and effect size and power estimates where appropriate.

## Author contributions

KML, LAP, CLW and MMM designed experiments. KML and MMM wrote the manuscript. KML performed pharmacology treatments, in vitro experiments, immunofluorescence and immunohistochemistry, histology, western blot, immunoassays, adult surgeries, behavioral experiments, cell counting, designed and made figures, and analyzed data. LAP performed pharmacological treatments, in vitro experiments, immunofluorescence and immunohistochemistry, histology, cell counting and reconstruction, biochemical measurements, analyzed data, and designed and made figures. CLW performed in vitro experiments, immunohistochemistry, counting and reconstruction, adult surgeries, behavioral experiments, biochemical measurements and analyzed data. KD assisted during the execution of *in vitro* experiments and biochemical measurements, performed histology, immunohistochemistry, and cell counting and reconstruction. AG generated allergic challenge animals, performed sex behavioral testing, olfactory investigation behavioral testing, immunohistochemistry and histology.

## Acknowledgements

We wish to acknowledge Shyama Patel from the Weill Cornell Medical College for assistance and training in brain mast cell isolations. We are also grateful to The Ohio State University Targeted Metabolomics Laboratory (metabolomics.osu.edu) for access to their LC-MS/MS equipment funded by the Translational Plant Sciences Targeted Investment in Excellence (TIE) and for performing LC-MS/MS analysis of histamine turnover in brain tissue. This work was supported by grant R01 MH052716 to MMM, F31NS093947 to LAP, and F32NS076327, R21MH105826 and The Ohio State University Startup Funds to KML. The authors have no conflicts of interest to declare.

---

**Figure 1, S1.**
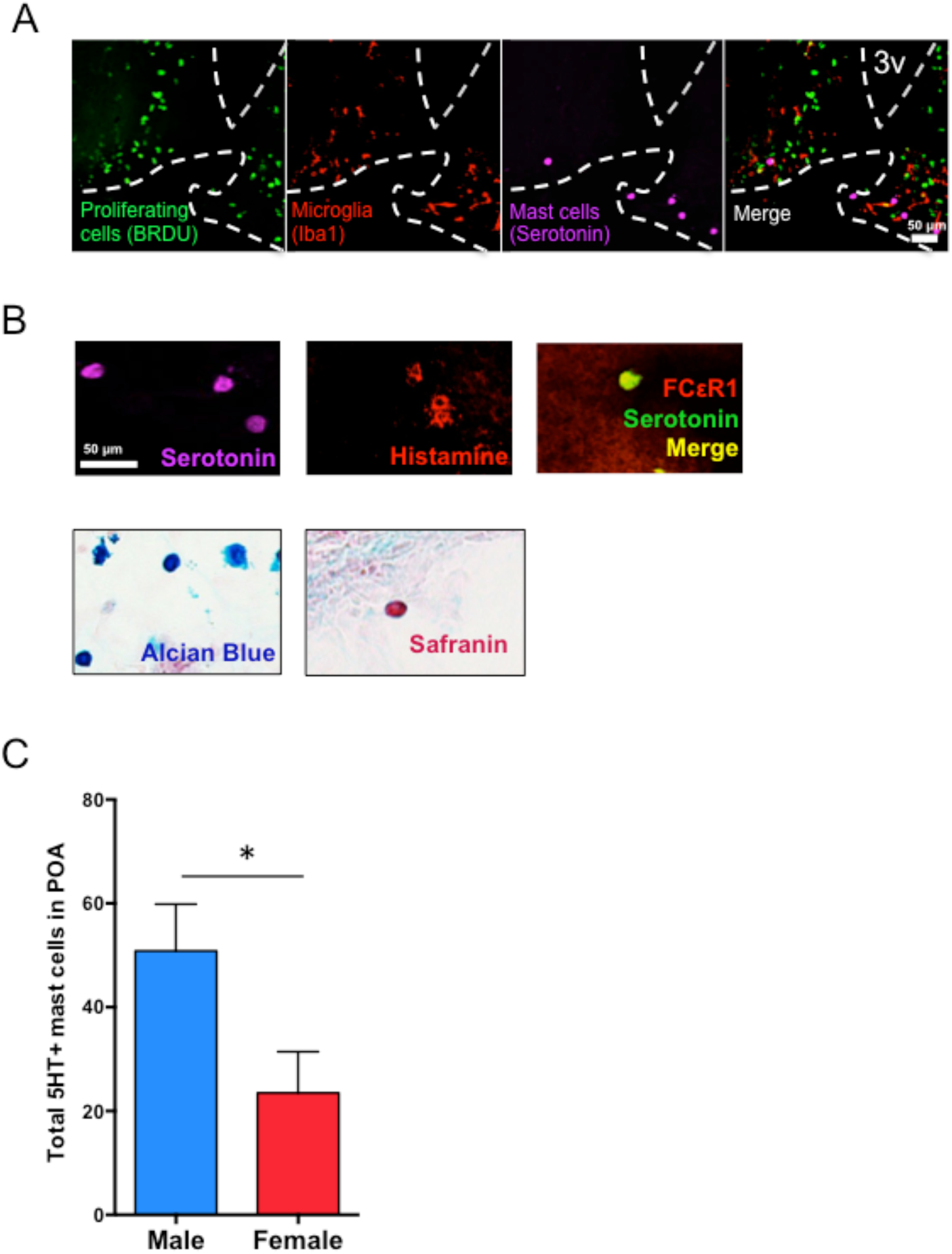
Mast cell proliferation and phenotype. (A) Labeling of proliferating cells (green, BrdU labeled), microglia (red, Iba1 labeled), mast cells (purple, serotonin labeled) and merged images (right) in the POA of the newborn animal on PN0 (dotted white lines indicate 3^rd^ ventricle and leptomeninges. There was no evidence of mast cell proliferation in the neonatal POA. (B) All mast cells in the POA stained by Alcian Blue (second from bottom) also immunolabel for serotonin (top), histamine (second from top) and the IgE receptor FC∊R1 (middle) indicating a mucosal phenotype. Occasional safranin-positive mast cells (bottom), indicative of a connective tissue phenotype, were detected in the meninges near the cortex, but not in the POA neuropil. (C). Counts of serotonin (5HT) positive mast cells in the POA on PN2 corroborated toluidine blue staining counts, with males having significantly more mast cells than females [t_(10)_=2.273, p = 0.046]. *indicates p<0.05. Group sizes: Panel C: ♂ n=6; ♀ n=6.

**Figure 1, S2.**
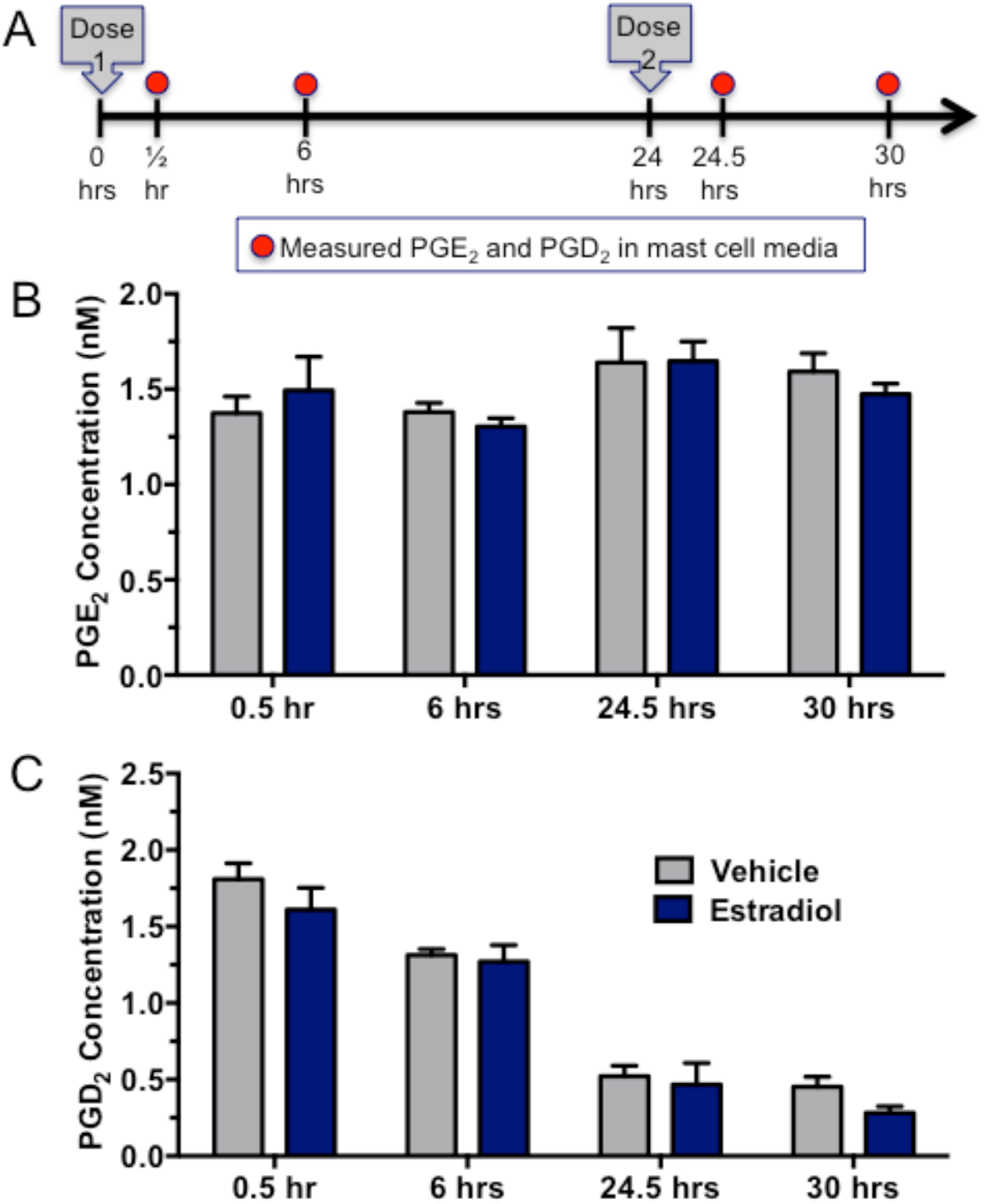
Mast cell prostaglandin production in response to estradiol. (A) Primary mast cells cultured from the brain were treated with estradiol and measured for concentration of PGE2 (B) and PGD2 (C) 0.5, 6, 24.5 and 30 hr after initial treatment, but the concentration of the prostaglandins did not increase at any time point (B: F_treat(1,16)_ = 0.04739, p = 0.8304); C: F_treat (1, 16)_=2.924, p = 0.107). Estradiol was given a second time, 24 hr after initial treatment, to the 24.5 and 30 hr samples and still did not evoke PGE2 or PGD2 release from mast cells. Group sizes: all groups: n=3.

**Figure 2, S1.**
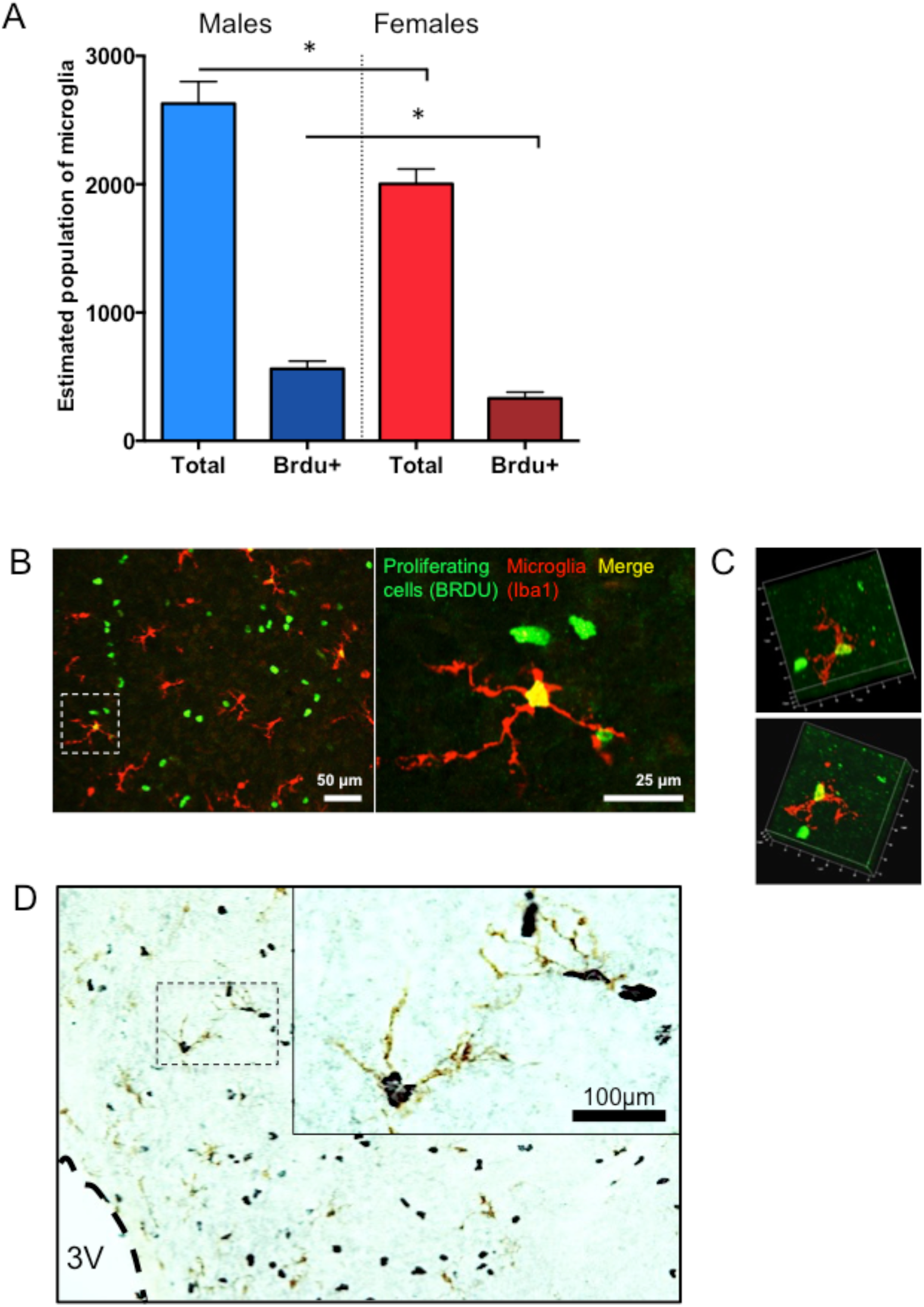
Sex differences in microglial proliferation in the developing POA. (A) Males had more microglia than females in the POA on PN2, [t_(17)_=2.78, *p*=0.01]. Unlike mast cells, Iba1-positive microglia co-localize with BRDU in the POA on PN2, indicating microglial proliferation, and males have higher numbers of proliferating microglia than females [t_(17)_=2.95, *p*=0.01]. (B) Representative images and (C) Representative Z-stack of immunofluorescently-stained BRDU-(green) and Iba1-(red) positive cells, showing co-localization. (D) Representative image of immunohistochemistry to double label microglia (Iba1 stained with DAB; brown) and proliferating cells (BRDU stained with Ni-DAB; black) in the POA on PN3 (3V = third ventricle). Counts of proliferating microglia were performed on DAB-stained tissue as seen in panel (D) and co-localization confirmed via confocal imaging (B-C). *indicates p<0.05. Group sizes: ♂ n=11; ♀ n=8.

**Figure 2, S2.**
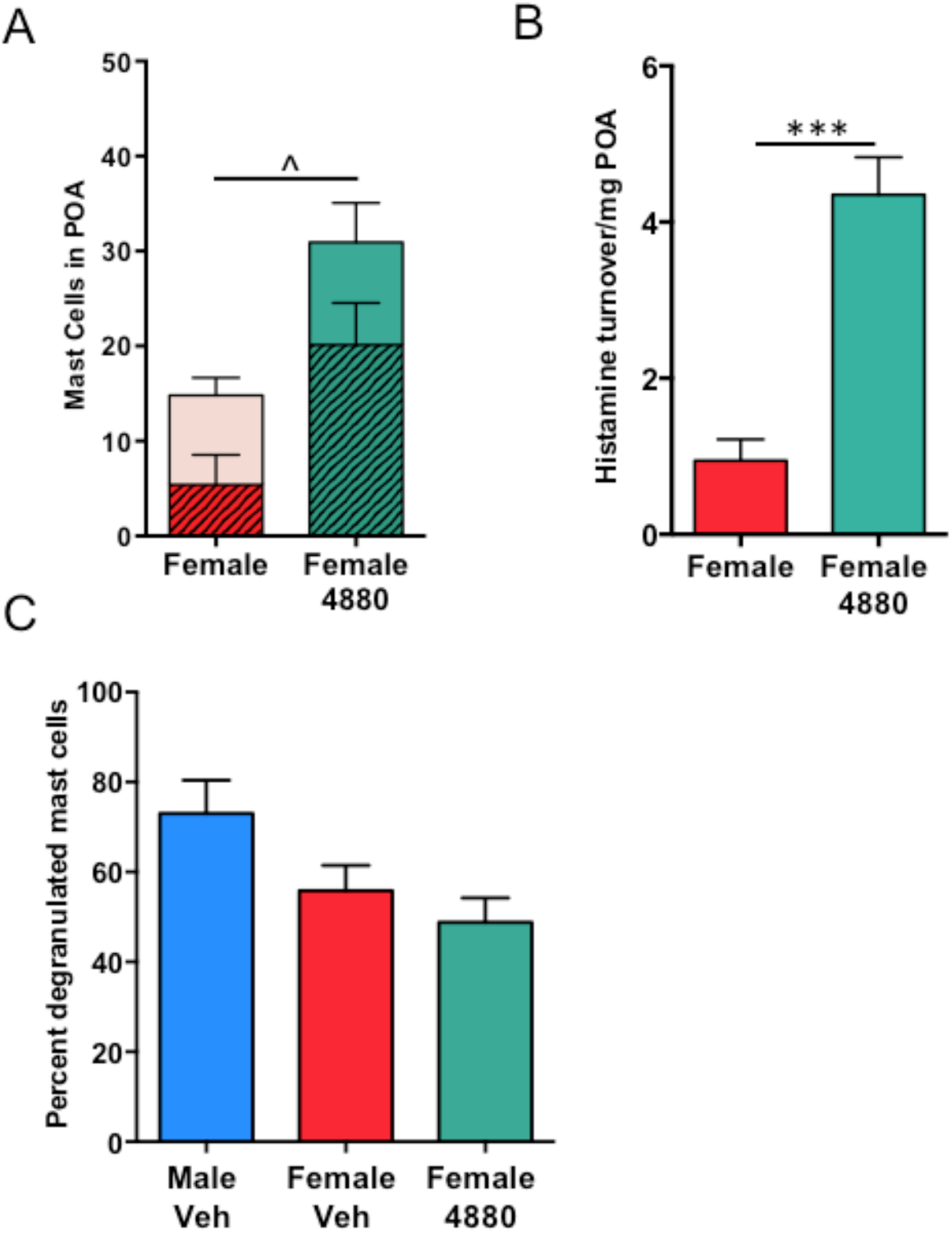
Confirmation of central versus peripheral mast cell manipulations. (A-B): Compound 48/80 treatment in vivo (icv) led to significant increases in (A) percentage of degranulated mast cells in the POA of females [t_(7)_=3.214, p=0.0148; full bars indicate total number of mast cells, shaded bars indicate number of degranulated mast cells] as well as (B) increased histamine turnover in the POA as measured by LC-MS/MS [t_(8)_=6.185, p=0.0003]. (C): Treatment with compound 48/80 did not change the proportion of degranulated mast cells in the spleen. ***indicates p<0.001, ^ indicates p<0.05 for % degranulated mast cells. Group sizes: A: ♀V: n=4; ♀ 4880 n=5. B: all groups n=5. C: ♂V n=5; ♀V n=3, ♀4880 n= 4.

**Figure 5, S1.**
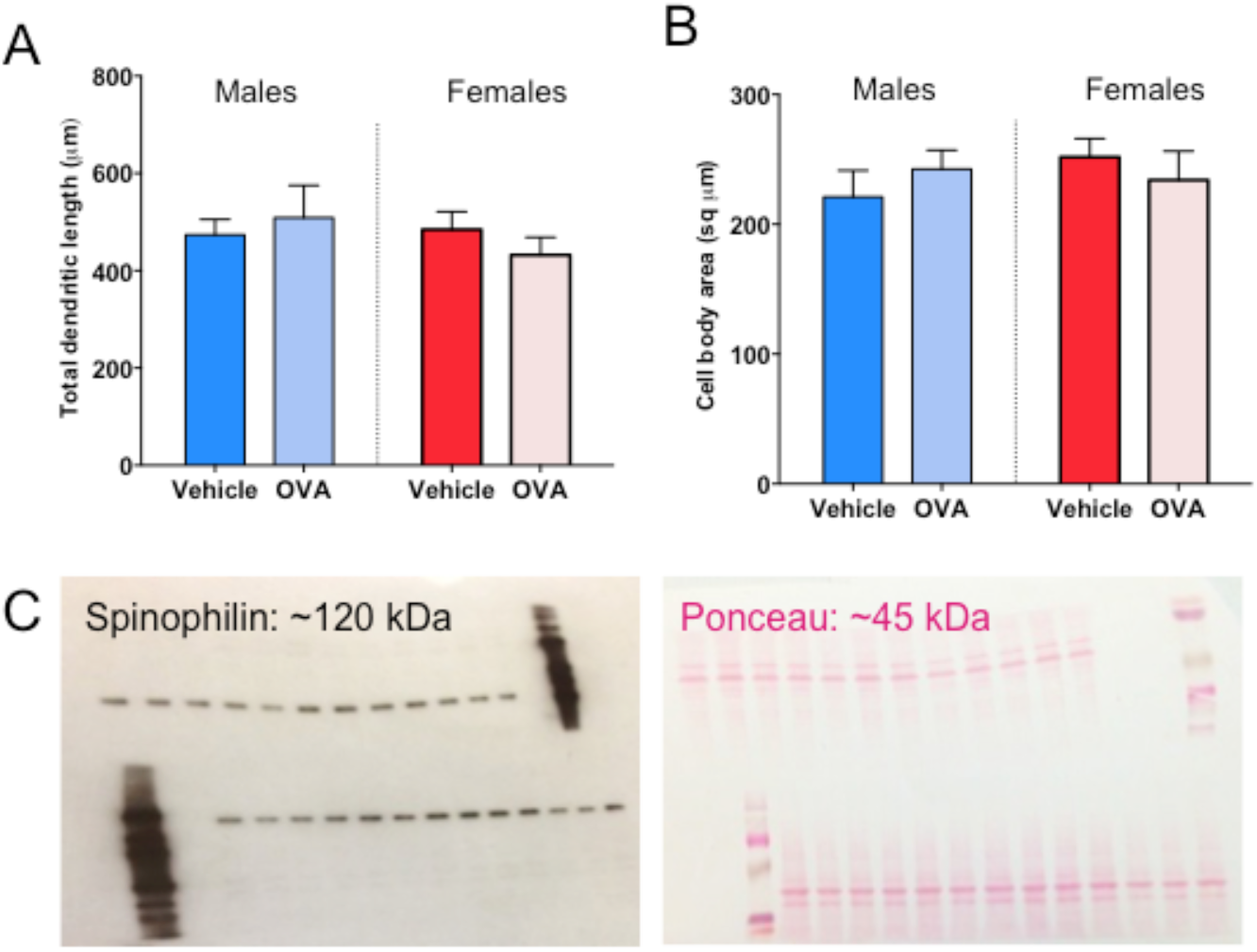
Mast cell effects on other neuronal morphology measures in POA. (A-B) While *in utero* allergic challenge affected the density of dendritic spines (see Fig. 5A-B), treatment did not influence the overall length of dendrites (A; F_int(1,12)_=0.94 p=0.351; F_treat(3,16)_=0.035, p=0.855; F_sex_=0.538 p=0.478) or the size of the soma (B; F _int(3,22)_=1.167 p=0.301; F_treat(3,16)_=0.10, p=0.921; F_sex_=0.379 p=0.550). (C) Full length western blot shown cropped in Fig. 4D: spinophilin (left; band at approximately 120 kDa) and Ponceau S stain of the same membrane (right; band at approximately 45 kDa). Group sizes for A-B: All groups: n=4.

**Fig. 6, S1.**
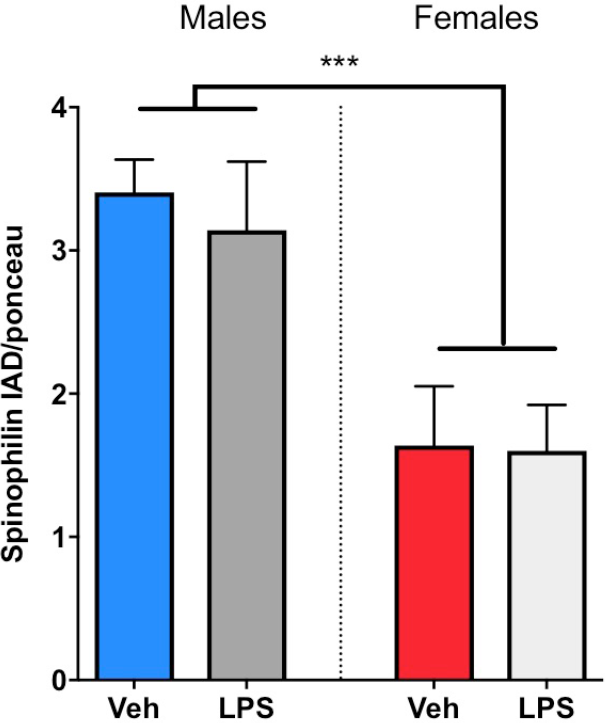
Effects of postnatal immune challenge on spinophilin content in the POA. Immune challenge with lipopolysaccharide (LPS) on the day of birth did not have a significant effect on spinophilin content in the POA of males or females on PN4 [F_(1,20)_=0.16, p=0.69]. There was a significant sex difference in spinophilin at PN4 [F_(1,20)_=18.62, p=0.0003], ***indicates p<0.001. Group sizes: ♂V n=5; ♂LPS n=5, ♀V n=7, ♀ LPS n=7.

**Fig. 6, S2.**
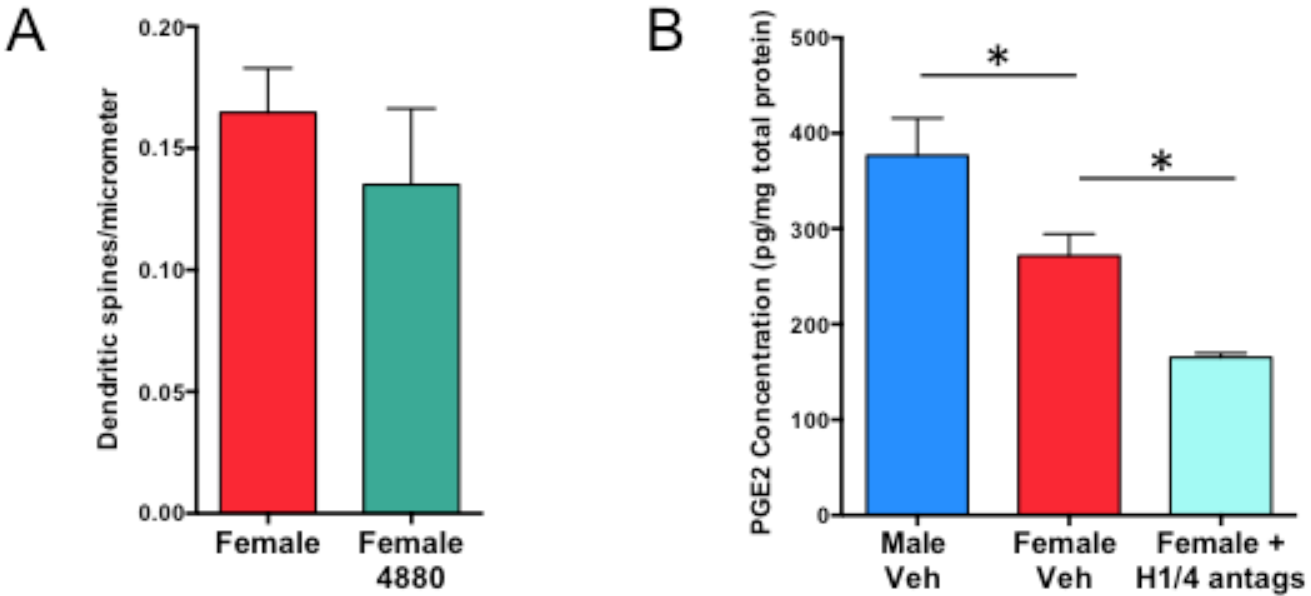
Direct effects of Compound 48/80 on primary neuronal cultures and PGE2 concentration in vivo in response to histamine receptor antagonism. (A) Compound 48/80 had no effects on the density of dendritic spines when given directly to the neurons [F_(2,77)_ = 2.07, p=0.13]. This contrasts with the spinogenic effects of Compound 48/80 when it is first administered to mast cells, and the mast cell conditioned media is then applied to the neurons (see Fig. 6A). (B) In vivo, the neonatal male POA showed higher PGE2 concentrations than females, and treating females with histamine receptor 1/4 antagonists further decreased PGE2 levels [F_(2,22)_=13.08 p=0.0002 HSD (M v F) p=0.0255; (F v F+H1/4ant) p=0.0.255]. *indicates p<0.05. Group sizes: A: All groups n=4. B: ♂V n=9; ♀V n=9, ♀H1/4 antags n= 7.

## References

1. E. M. Hull, R. L. Meisel, B. D. Sachs. Male sexual behavior. Horm. brain Behav. 1, 3–137 (2002).

2. M. V., Wu, D. S. Manoli, E. J. Fraser, J. K. Coats, J. Tollkuhn, S. Honda, N. Harada, N. M. Shah. Estrogen masculinizes neural pathways and sex-specific behaviors. Cell 139, 61–72. (2009).

3. M. M. McCarthy. Estradiol and the developing brain. Physiol. Rev. 88, 91–124 (2008).

4. S. K. Amateau, M. M. McCarthy. Induction of PGE2 by estradiol mediates developmental masculinization of sex behavior. Nat. Neurosci. 7, 643–50 (2004).

5. K. M. Lenz, B. M. Nugent, R. Haliyur, M. M. McCarthy. Microglia are essential to masculinization of brain and behavior. J. Neurosci. 33, 2761–72 (2013).

6. J. W. VanRyzin, S. J. Yu, M. Perez-Pouchoulen, M.M. McCarthy. Temporary depletion of microglia during the early postnatal period induces lasting sex-dependent and sex-independent effects on behavior in rats. eNeuro.0297-16.2016 (2016).

7. R. Silver, J. P. Curley, Mast cells on the mind: new insights and opportunities. Trends Neurosci. 36, 513–21 (2013).

8. S. K. Amateau, J. J. Alt, C. L. Stamps, M. M. McCarthy. Brain estradiol content in newborn rats: sex differences, regional heterogeneity, and possible de novo synthesis by the female telencephalon. Endocrinology. 145, 2906–17 (2004).

9. F. Aloisi. Immune function of microglia. Glia. 179, 165–179 (2001).

10. Y. Wu, L. Dissing-Olesen, B. A. MacVicar, B. Stevens. Microglia: Dynamic Mediators of Synapse Development and Plasticity. Trends Immunol. 36, 605–13 (2015).

11. K. Nautiyal, C. A. Dailey, J. L. Jahn, E. Rodriguez, N. H. Son, J. V. Sweedler, R. Silver. Serotonin of mast cell origin contributes to hippocampal function. Eur. J. Neurosci. 36, 2347–2359 (2012).

12. D. Chatterjea, A. Wetzel, M. Mack, C. Engblom, J. Allen, C. Mora-Solano, L. Paredes, E. Balsells, T. Martinov. Mast cell degranulation mediates compound 48/80-induced hyperalgesia in mice. Biochem. Biophys. Res. Commun. 425: 237–43 (2012).

13. S. Chikahisa, T. Kodama, A. Soya, Y. Sagawa, Y. Ishimaru, H. Sei, S. Nishino. Histamine from brain resident MAST cells promotes wakefulness and modulates behavioral states. PLoS One 8(10):e78434 (2013).

14. K. M. Lenz, C. L. Wright, R. C. Martin, M. M. McCarthy. Prostaglandin E_2_ regulates AMPA receptor phosphorylation and promotes membrane insertion in preoptic area neurons and glia during sexual differentiation. PLoS One. 6, e18500 (2011).

15. C. L. Wright, M. M. McCarthy. Prostaglandin E2-induced masculinization of brain and behavior requires protein kinase A, AMPA/kainate, and metabotropic glutamate receptor signaling. J. Neurosci. 29, 13274–82 (2009).

16. M.J. Baum, B.J. Everitt. Increased expression of c-fos in the medial preoptic area after mating in male rats: role of afferent inputs from the medial amygdala and midbrain central tegmental field. Neuroscience 50, 627–46. (1992).

17. JM Dominguez, H. Gil, E.M. Hull. Preoptic glutamate facilitates male sexual behavior. J Neurosci 26, 1699–1703. (2006).

18. S. Georgin-Lavialle, D. S. Moura, A. Salvador, J.C. Chauvet-Gelinier, J. M. Launay, G. Jamaj, F. Cote, E. Soucle, M. O. Chandesris, S. Barete, C. Grandpeix-Guyodo, C. Bachmeyer, M. A. Alvanakian, A. Aouba, O. Lortholary, P. Dubreuil, J. R. Teyssier, B. Trojak, E. Haffen, P. Vandel, B. Bonin, French Mast Cell Study Group, O. Hermine, R. Gaillard. Mast cells’ involvement in inflammation pathways linked to depression: evidence in mastocytosis. Mol. Psychiatry (2016), doi:10.1038/mp.2015.216.

19. K. J. Zucker, S. J. Bradley. Gender Identity Disorder and Psychosexual Problems in Children and Adolescents (Guilford, New York, 1995).

20. M. Hines. Gender development and the human brain. Annu. Rev. Neurosci. 34, 69–88 (2011).

21. E. Mackey, S. Ayyadurai, C.S. Pohl, S. D’Costa, Y. Li, A.J. Moser. Sexual dimorphism in the mast cell transcriptome and the pathophysiological responses to immunological and psychological stress. Biol. Sex Diff. 7:60 doi: 10.1186/s13293-016-0113-7 (2016).

22. D. M. Werling, N. N. Parikshak, D. H. Geschwind. Gene expression in human brain implicates sexually dimorphic pathways in autism spectrum disorders. Nat. Commun. 7, 10717 (2016).

23. J. M. Schwarz, S. D. Bilbo. Sex, glia, and development: interactions in health and disease. Horm. Behav. 62:243–53. (2012).

24. J. T. Instanes, A. Halmoy, A. Engeland, J. Haavik, K. Furu, K. Klungsoyr. Attention-Deficit/Hyperactivity Disorder in Offspring of Mothers With Inflammatory and Immune System Diseases. Biol. Psychiatry doi:10.1016/j.biopsych.2015.11.024 (2015).

25. M. L. Estes, A. K. McAllister. Immune mediators in the brain and peripheral tissues in autism spectrum disorder. Nat. Rev. Neurosci. 16, 469–486 (2015).

26. K. Suzuki, G. Sugihara, Y. Ouchi, K. Nakamura, M. Futasubashi, K. Takebayashi, Y, Yoshihra, K. Omata, K. Matsumoto, K.J. Tsuchiya, Y. Iwata, M. Tsuji, T. Sugiyama, N. Mori. Microglial activation in young adults with autism spectrum disorder. JAMA psychiatry. 70, 49–58 (2013).

27. D. Braunschweig, J. Van de Water. Maternal autoantibodies in autism. Arch. Neurol. 69, 693–9 (2012).

28. W. Mandy, R. Chilvers, U. Chowdhury, G. Salter, A. Seigal, D. Skuse. Sex differences in autism spectrum disorder: evidence from a large sample of children and adolescents. J. Autism Dev. Disord. 42, 1304–13 (2012).

29. S. M. Schaafsma, D. W. Pfaff. Etiologies underlying sex differences in Autism Spectrum Disorders. Front. Neuroendocrinol. 35, 255–71 (2014).

30. K. M. Abel, R. Drake, J. M. Goldstein. Sex differences in schizophrenia. Int. Rev. Psychiatry. 22, 417–28 (2010).

31. J. M. Goldstein, L. J. Seidman, S. Santangelo, P. H. Knapp, M. T. Tsuang. Are schizophrenic men at higher risk for developmental deficits than schizophrenic women? Implications for adult neuropsychological functions. J. Psychiatr. Res. 28, 483–498 (1994).

32. P. Nopoulos, M. Flaum, N. C. Andreasen. Sex Differences in Brain Morphology in Schizophrenia. Am. J. Psychiatry. 154, 1648–1654 (1997).

33. T. R. Insel, L. J. Young. The neurobiology of attachment. Nat. Rev. Neurosci. 2, 129–36 (2001).

34. J. N. Ferguson, J. M. Aldag, T. R. Insel, L. J. Young. Oxytocin in the Medial Amygdala is Essential for Social Recognition in the Mouse. J. Neurosci. 21, 8278–8285 (2001).

35. F. A. Champagne, I. C. Weaver, J. Diorio, S. Dymov, M. Szyf, M. J. Meaney. Maternal care associated with methylation of the estrogen receptor-alpha1b promoter and estrogen receptor-alpha expression in the medial preoptic area of female offspring. Endocrinology. 147, 2909–15 (2006).

36. T. C. Theoharides, A. Angelidou, K. D. Alysandratos, B. Zhang, S. Asadi, K. Francis, E. Toniato, D. Kalogeromitros. Mast cell activation and autism. Biochim. Biophys. Acta. 1822, 34–41 (2012).

37. S. D. Patel, G. Brennan, J. Brazin, A. J. Ciardiello, R. B. Silver, S. J. Vannucci. Mast cell isolation from the immature rat brain. Dev. Neurosci. 35, 265–71 (2013).

38. G. Krishnaswamy, D. S. Chi. Mast Cells: Methods and Protocols (Humana Press, Totowa, NJ, 2006).

39. E. K. Murray, M. M. Varnum, J. L. Fernandez, G. J. de Vries, N. G. Forger. Effects of neonatal treatment with valproic acid on vasopressin immunoreactivity and olfactory behavior in mice. J. Neuroendocrinol. 23, 906–914.

